# History-Dependent Physiological Adaptation to Lethal Genetic Modification under Antibiotic Exposure

**DOI:** 10.1101/2021.09.05.459045

**Authors:** Yuta Koganezawa, Miki Umetani, Moritoshi Sato, Yuichi Wakamoto

## Abstract

Genetic modifications, such as gene deletion and mutations, could lead to significant changes in physiological states or even cell death. Bacterial cells can adapt to diverse external stresses, such as antibiotic exposure, but can they also adapt to detrimental genetic modification? To address this issue, we visualized the response of individual *Escherichia coli* cells to deletion of the antibiotic resistance gene under chloramphenicol (Cp) exposure, combining the light-inducible genetic recombination and microfluidic long-term single-cell tracking. We found that a significant fraction (∼ 40%) of resistance-gene-deleted cells demonstrated a gradual restoration of growth and stably proliferated under continuous Cp exposure without additional mutations. Such physiological adaptation to genetic modification was not observed when the deletion was introduced in 10 h or more advance before Cp exposure. Resistance gene deletion under Cp exposure disrupted the stoichiometric balance of ribosomal large and small subunit proteins (RplS and RpsB). However, the balance was gradually recovered in the cell lineages with restored growth. These results demonstrate that bacterial cells can adapt even to lethal genetic modifications by plastically gaining physiological resistance. However, the access to the resistance states is limited by the environmental histories and the timings of genetic modification.

## Introduction

Bacteria in nature are constantly challenged by environmental and genetic perturbations and must be robust against them for survival. Bacterial cells possess programs to adapt or resist such perturbations. For example, under the conditions of nutrient deprivation, *E. coli* and related bacteria provoke RpoS-mediated general stress response and globally change the metabolic and gene expression profiles to protect themselves from the stress [1]. DNA damage also induces SOS response, which promotes both DNA repair and mutagenesis [2, 3]. If the mutations in essential genetic elements remain unrepaired, their influence will be propagated to cellular phenotypes or even cause cell death. How rapidly new genetic changes alter cellular phenotypes and whether they always give rise to the same phenotypes in given environments are fundamental questions in genetics [4, 5]. Nevertheless, such time-dependent and redundant genotype-phenotype correspondences are usually deemed negligible or insignificant in most genetics analyses.

However, the evidence suggests that biological systems can buffer or compensate for the impact of genetic changes at the cytoplasmic and physiological levels [6]. For example, specific molecular chaperons could hinder genetic variations from manifesting as morphological and growth phenotypes in fruit flies [7], plants [8], and bacteria [9]. Furthermore, the loss of biological functions caused by genetic changes can be compensated for by modulating the RNA and protein levels of the mutated or other related genes in many organisms [10]. In *Bacillus subtilis*, the expression noise of mutant sporulation regulator results in the partial penetrance of its influence to spore-forming phenotypes [11]. These observations across diverse organisms suggest that phenotypic consequences of genetic modifications can be modulated based on environmental and physiological contexts, which may promote the survival and evolution of the organisms.

Despite these experimental implications, it remains elusive whether bacterial cells can circumvent even lethal genetic modifications such as antibiotic resistance gene deletion under antibiotic exposure. Furthermore, how cells initially respond to the genetic changes and how their physiological and phenotypic states are modulated in longer timescales are poorly characterized. However, addressing these issues requires precisely defining the timings of genetic changes and tracking individual cells for a long period to unravel their phenotypic transitions and consequences [5].

In this study, we resolve these technical issues by combining the photoactivatable Cre (PA-Cre) light-inducible genetic recombination technique [12] and microfluidic long-term single-cell tracking [13]. We induced a pre-designed deletion of chromosomally-encoded and fluorescently-tagged drug resistance gene in *E. coli* directly in the microfluidic device. The results show that all of the resistance-gene-deleted cells under continuous drug exposure showed a decline in growth in 5-7 generations, but a fraction of the resistance-gene-deleted cells gradually restored their growth without additional mutations. In contrast, no cells restored growth when the same deletion was introduced 10 h or more in advance before drug exposure. Therefore, bacterial cells can physiologically adapt to lethal genetic modifications. However, its feasibility depends on environmental histories and the timings of genetic modifications.

## Results

### Resistance gene deletion in *E. coli* by blue-light illumination

To investigate the response of individual cells to antibiotic resistance gene deletion, we constructed an *E. coli* strain expressing chloramphenicol acetyltransferase (CAT) tagged with mCherry red fluorescent protein (Fig. 1A). CAT confers resistance to chloramphenicol (Cp) by acetylating Cp [14]. The *mcherry-cat* resistance gene was integrated on the chromosome along with the upstream and downstream *loxP* sequences so that the resistance gene could be excised by Cre recombinase (Fig. 1A). We also introduced the PA-Cre recombination system, which consists of split Cre-recombinase fragments, called CreC and CreN, attached to p-Magnet (p-Mag) and n-Magnet (n-Mag) monomers, respectively (Fig. 1B) [15]. p-Mag and n-Mag are heterodimers derived from the fungal photo-receptor, Vivid [15], which heterodimerize upon blue-light illumination. In the PA-Cre system, blue-light illumination leads to heterodimerizations of p-Mag-CreC and n-Mag-CreN fragments and recovers Cre recombination activity (Fig. 1B) [12]. Therefore, this system allows us to induce the resistance gene deletion at arbitrary timings by blue-light illumination. The original PA-Cre system was designed for use in mammalian cells [12]; we thus replaced the plasmid backbone and the promoter for use in *E. coli*.

**Figure 1.**
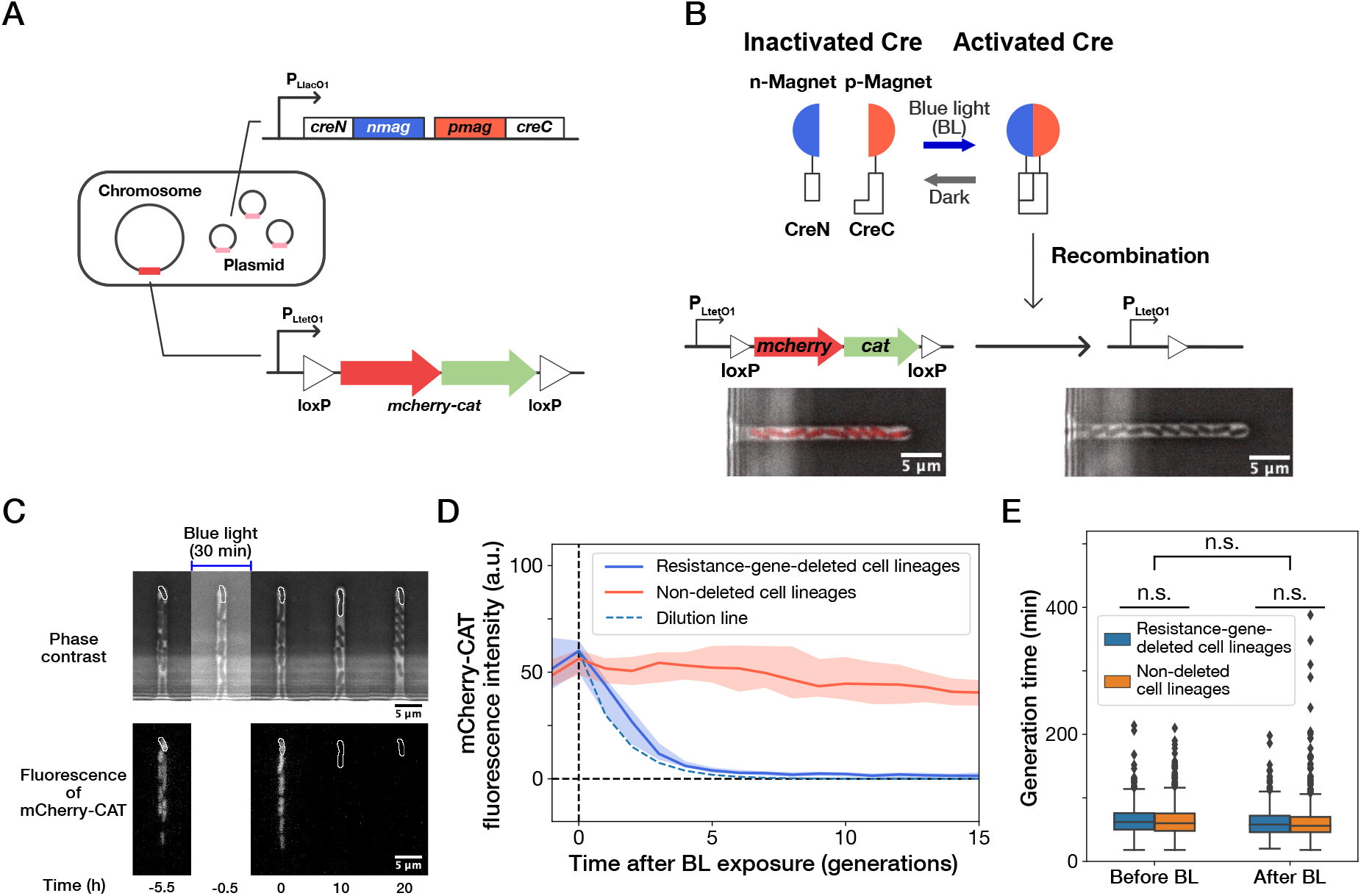
Live-cell monitoring of phenotypic transitions in response to gene deletion. (A) Schematic drawing of an *E. coli* strain, YK0083. This strain harbors the photo-removable *mcherry-cat* gene on the chromosome and a low-copy plasmid carrying the pa-cre genes (*creN-nmag* and *pmag-creC*). (B) The PA-Cre system. Blue-light illumination provokes the dimerization of two PA-Cre fragments and induces the deletion of the *mcherry-cat* gene. The micrographs represent the combined images of phase contrast and mCherry-CAT fluorescence channels. Left and right images show the cells before and after blue-light illumination, respectively. (C) Representative time-lapse images of gene-deletion experiments. The upper and lower images show the cells in phase contrast and mCherry-CAT fluorescence channels, respectively. The cell lineages at the closed end of the growth channel (outlined in white) were monitored. A 30-min blue-light illumination starting at *t* = −0.5 h led to the loss of mCherry-CAT fluorescence signals in this cell lineage. (D) The transitions of mCherry-CAT fluorescence intensities in resistance-gene-deleted (blue) and non-deleted (red) cell lineages. Lines and shaded areas represent the medians and the 25-75% ranges, respectively. The cyan broken line represents the expected fluorescence decay curve when the fluorescence intensity decreases to half in each generation. (E) Generation time of resistance-gene-deleted and non-deleted cells ten generations before and after blue-light illumination. The middle line and both edges of the boxes represent the medians and the 25-75% ranges of generation time. Whiskers indicate the minimum and maximum of the data except for the outliers. The points represent the outliers. No significant differences in generation time were detected between the groups at the significance level of 0.01 (*p* = 0.47 for before BL, *p* = 0.027 for after BL, *p* = 0.055 for before BL vs after BL, Mann-Whitney U test).

We first analyzed the response of the constructed strain YK0083 to blue-light illumination under drug-free conditions. Single-cell observations with the mother machine microfluidic device [13] and custom stage-top LED illuminator (Fig. S1) revealed that 30-min blue-light exposure (*λ* =464 ∼ 474 nm, 6.8 mW at the specimen position) led to the loss of mCherry-CAT fluorescence in 25% (50/200) cells (Fig. 1C, D, and Movie S1), suggesting the deletion of the *mcherry-cat* gene in these cell lineages. The fluorescence signals decayed to the background level in 4-5 generations, and the decay kinetics was consistent with the dilution by growth (Fig. 1D). Furthermore, the 30-min blue-light illumination and genetic recombination did not affect the growth of individual cells (Fig. 1E).

Extending illumination duration increased the frequency of cells showing loss of fluorescence, reaching 100% (*n* = 296) in 4 h (Fig. S2A and B). We confirmed the correspondence between the loss of mCherry fluorescence and *cat* gene deletion by colony PCR (Fig. S2C).

### Fractional continuation of growth against resistance gene deletion under drug exposure

We next induced the deletion of the resistance gene in YK0083 cells in the mother machine, continuously flowing a medium containing 15 *µ*g/ml of Cp. This drug concentration was 1.5-fold higher than the minimum inhibitory concentration (MIC) of the non-resistant strain YK0085 (10 *µ*g/ml), which was constructed by illuminating the YK0083 cells by blue light in batch culture (Fig. S3). Therefore, it was expected that this drug concentration would inhibit the growth of resistance-gene-deleted cells. This Cp concentration was significantly lower than the MIC of resistant YK0083 cells (100 *µ*g/ml, Fig. S3) and did not influence their growth rates (Fig. S4).

A 30-min blue-light illumination induced the *mcherry-cat* gene deletion in 24.5% (343/1399) in YK0083 cells (Fig. 2A and Movie S2). While non-deleted cells continued to grow with their generation time (interdivision time) and mCherry-CAT fluorescence intensity nearly unaffected (Fig. 2B and C), resistance-gene-deleted cells gradually showed a decline in growth (Fig. 2D-G). The fluorescence intensity of the resistance-gene-deleted cells decayed to the background levels in 4-5 generations (Fig. 2D, F, and H), and their generation time also increased correspondingly (Fig. 2E, G, and I). However, while the growth of 62.7% (163/260) of resistance-gene-deleted cells was eventually stopped, the other 37.3% cells restored and continued their growth over 30 generations without the *cat* resistance gene (Fig. 2H and I). The generation time of these cell lineages recovered from 6.3 h (the median of 6th-9th generations) to 3.0 h (the median of 21st-30th generations) under the continuous drug exposure (Fig. 2I). The elongation rate transitions across multiple generations also manifested the growth recovery (Fig. S5).

**Figure 2.**
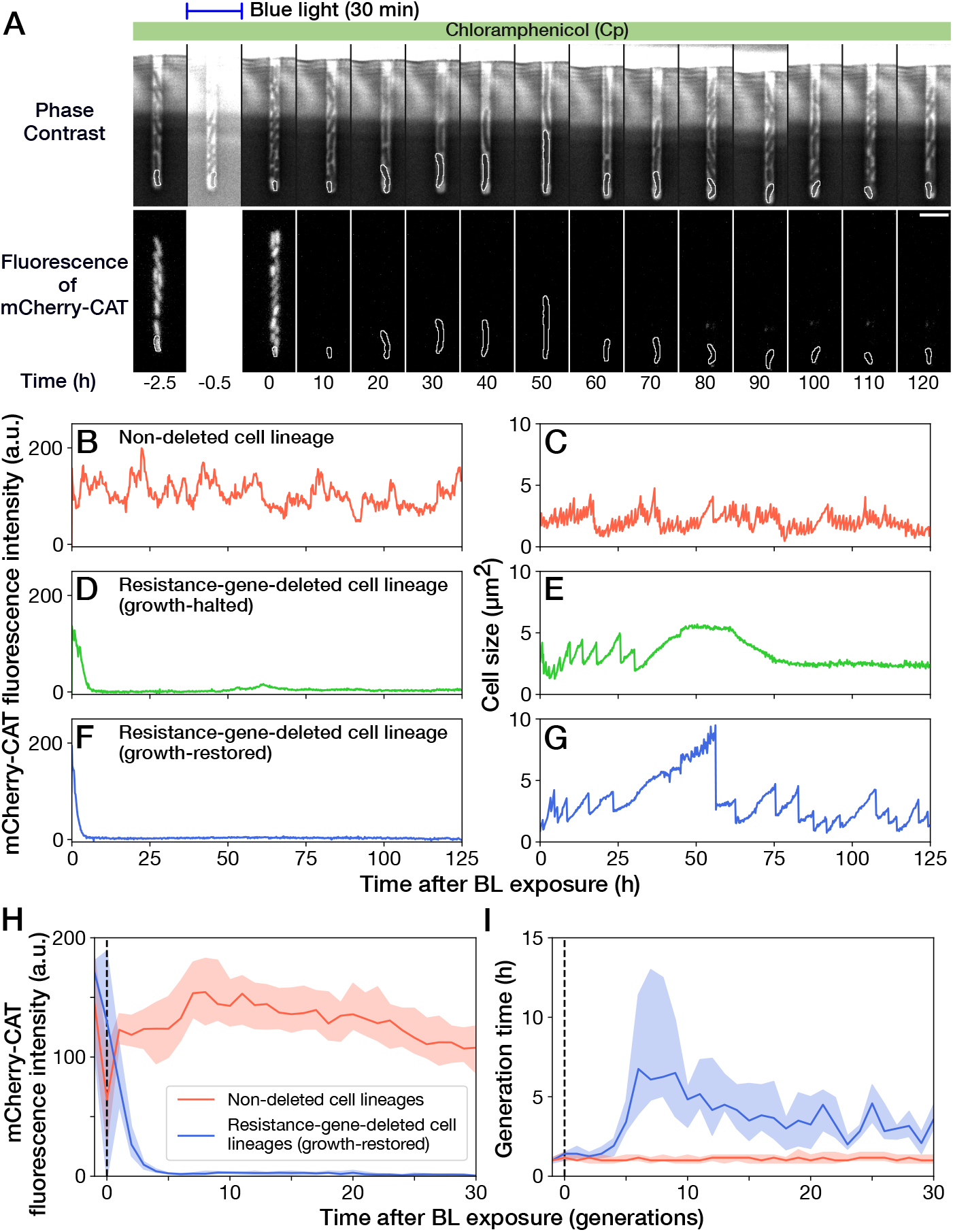
Growth continuation under Cp exposure against *cat* gene deletion. (A) Time-lapse images of a cell lineage that continued growth and division against *mcherry-cat* gene deletion. The upper and lower sequences show phase contrast and mCherry fluorescence images, respectively. Blue light was illuminated from *t* = −0.5 h to 0 h. The cells at the closed end of the growth channel were monitored during the experiment, which are outlined in white on the images. Scale bar, 5 *µ*m. (B-G) The transitions of mCherry-CAT fluorescence intensities and cell size in single-cell lineages. (B, C) Non-deleted cell lineage. (D, E) Growth-halted resistance-gene-deleted cell lineage. The decrease in cell size after 60 h was due to the shrinkage of the cell body. (F, G) Growth-restored resistance-gene-deleted cell lineage. (H, I) The transitions of mCherry-CAT fluorescence intensities (H) and generation time (I). The lines and shaded areas represent the medians and the 25-75% ranges, respectively. Red represents non-deleted cell lineages. Blue represents growth-restored resistance-gene-deleted cell lineages. The transitions are shown in generations.

We first suspected that the growth recovery under Cp exposure was attributed to the presence of the *cat* gene untagged from the *mcherry* gene or by additional unintended resistance-enhancing mutations. To test this hypothesis, we sampled the culture media flowing out from the microfluidic device and obtained cell populations derived from a single or few ancestral cells by limiting dilution (Fig. S6A). PCR analysis confirmed the lack of *cat* gene in the non-fluorescent cell populations (Fig. S6B). Furthermore, whole-genome sequencing of the cell populations obtained by limiting dilution showed no additional mutations in 4 out of 5 non-fluorescent cell populations tested (Table S1). One point mutation was present in one cellular population, but this mutation did not affect the MIC (Fig. S6C). Therefore, it was unlikely that unintended genetic changes were responsible for the growth restoration. It is also unlikely that the growth restoration was caused by the residual mCherry-CAT proteins because 30 consecutive cell divisions dilute cytoplasmic proteins 2^30^ ≈ 10^9^ folds if additional production is prevented; note that even the total number of protein molecules in a bacterial cell are in the order of 10^6^-10^7^ [16];

We also detected no significant differences in the mCherry-CAT fluorescence intensity before blue-light illumination between the growth-halted and growth-restored cell lineages (Fig. S7A). Furthermore, no significant differences were observed even between the resistance-gene-deleted and non-deleted cell lineages (Fig. S7A). We also examined the influence of elongation rate before blue-light illumination on the gene deletion and the fates after gene deletion, finding no correlations (Fig. S7B). Therefore, neither the amount of mCherry-CAT proteins at the time of gene deletion nor pre-deletion elongation rate affected the likelihood of gene deletion and the determination of growth-halt and growth-restoration fates under these experimental conditions.

### Resistance gene deletion long before Cp exposure prevents growth restoration

The high frequency of growth restoration observed against the resistance gene deletion was unexpected since the MIC of the susceptible strain was below 15 *µ*g/ml (Fig. S3 and S6C). To understand this discrepancy, we cultured the susceptible YK0085 cells in the mother machine first without Cp and then exposed them to 15 *µ*g/ml of Cp directly in the device. In contrast to the previous observation, all cells stopped growth and division entirely, with no cells restoring growth (“Pre-deleted” in Fig. 3A-C and Movie S3). This result excludes the hypothesis that the unique cultivation environments in the microfluidics device are the cause of growth restoration. Instead, this observation implies that the timing of gene deletion is crucial for the cells to withstand the Cp exposure without the resistance gene and restore growth.

**Figure 3.**
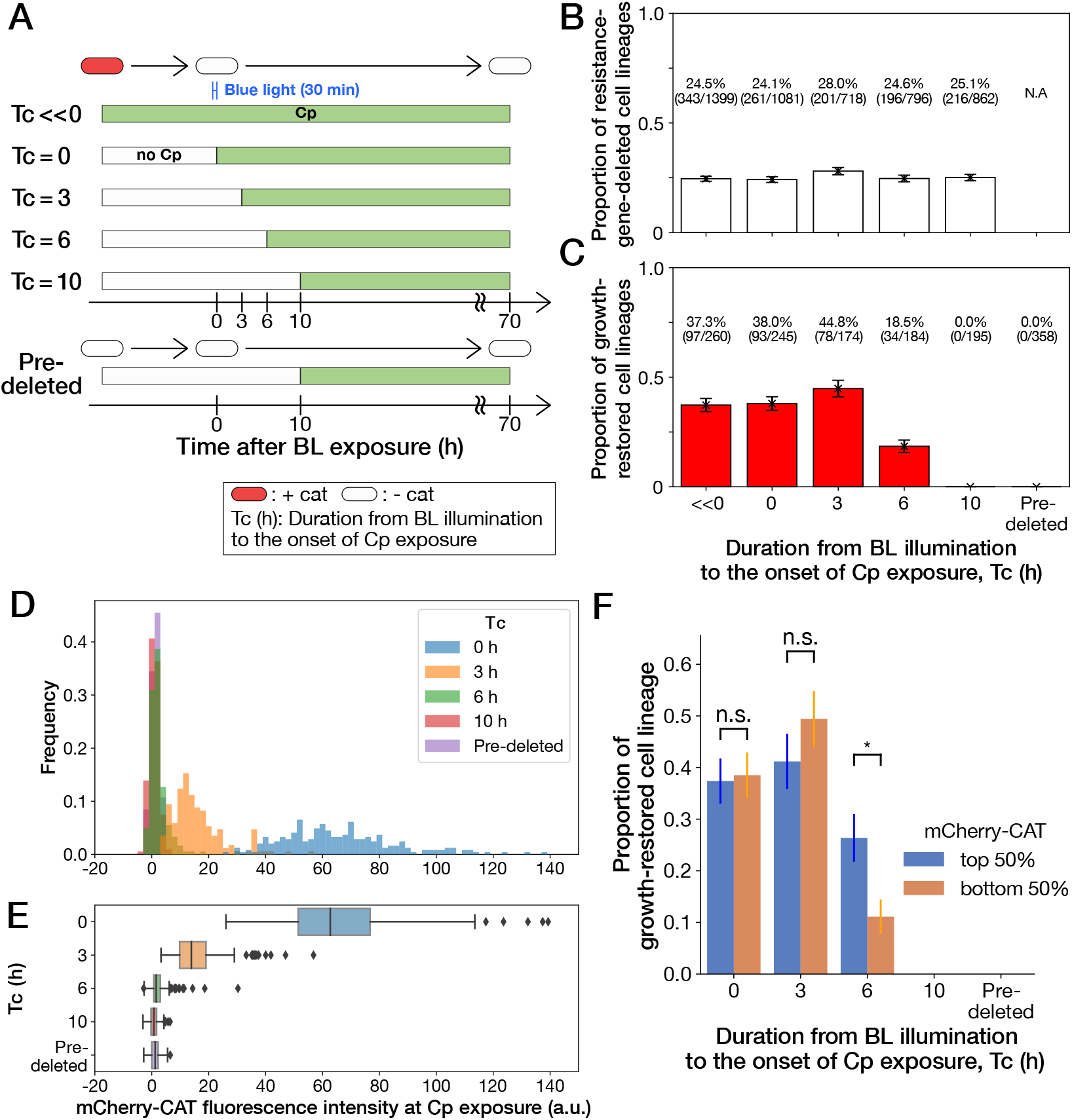
History-dependent maintenance of Cp resistance. (A) The schematic diagram of experiments. The duration from the end of blue-light illumination to the onset of Cp exposure (*T_c_*) was varied from 0 to 10 h. *T_c_* ≪ 0 represents the continuous Cp-exposure condition. “Pre-deleted” denotes the results of the experiments performed with the *mcherry-cat*-deleted YK0085 strain. The non-resistant YK0085 cells were not exposed to blue light. (B) Fractions of *mcherry-cat*-deleted cell lineages across different *T_c_* conditions. The numbers of resistance-gene-deleted cell lineages among the total numbers of cell lineages observed during the measurements are shown above the bars. Error bars represent standard errors. (C) Fractions of growth-restored cell lineages among resistance-gene-deleted cell lineages. The numbers of growth-restored cell lineages among all the resistance-gene-deleted cell lineages are shown above the bars. Error bars represent standard errors. The numbers of resistance-gene-deleted cell lineages are different from those in B because some cell lineages were flushed away from the growth channels at the later time points of the measurements, and their fates could not be determined. (D, E) The distributions of the mCherry-CAT fluorescence intensities at the onset of Cp exposure across the different *T_c_* conditions represented by histograms (D) and box plots (E). (F) Fractions of growth-restored cell lineages and their dependence on the mCherry-CAT fluorescence at the onset of Cp exposure. Blue and orange bars indicate the fractions of growth-restored cells among cell lineages whose mCherry-CAT fluorescence intensities were higher and lower than the median, respectively. Error bars represent standard errors. The growth-restored cell lineages were not detected under the *T_c_* = 10 h and “Pre-deleted” conditions. Fractional differences between the top 50% and the bottom 50% of cell lineages were statistically significant only for the *T_c_* = 6 h condition (*p* = 0.86 for *T_c_* = 0 h; *p* = 0.28 for *T_c_* = 3 h; and *p* = 8.6 × 10^−3^ for *T_c_* = 6 h, two proportional z-test).

To further investigate the importance of the timing of gene deletion, we performed microfluidic single-cell measurements with the resistant YK0083 strain, varying the duration from blue-light illumination to the onset of Cp exposure (*T*_*c*_) between 0 h to 10 h (Fig. 3A). The proportions of cells that lost the *mcherry-cat* gene in response to blue-light illumination almost remained unchanged among all conditions, ranging from 24-28% (Fig. 3B). However, the proportions of growth-restored cell lineages among resistance-gene-deleted cells strongly depended on *T*_*c*_ (Fig. 3C). When Cp exposure was initiated immediately after the blue-light illumination (*T*_*c*_ = 0 h) or after 3 h (*T*_*c*_ = 3 h), the proportions of growth-restored cells were nearly equivalent to those observed under continuous Cp exposure conditions (Fig. 3C). In contrast, the proportions of growth-restored cells were reduced to 18.5% (34/184) when *T*_*c*_ = 6 h, and we did not detect any growth restoration when *T*_*c*_ = 10 h (0/195; Fig. 3C). The almost equivalent frequencies of growth-restored cell lineages under the *T*_*c*_ = 0 h and *T*_*c*_ = 3 h conditions and those observed under continuous exposure exclude the possibility that growth restoration requires prior exposure to Cp before resistance gene deletion.

We next conjectured that a low amount of mCherry-CAT proteins is required at the onset of Cp exposure to withstand and restore growth without the resistance gene. In fact, *mcherry-cat* gene deletion before Cp exposure led to the dilution of mCherry-CAT proteins by growth by the time of Cp exposure (Fig. 3D and E).

To examine whether the mCherry-CAT concentration in individual cells affects growth restoration, we divided the resistance-gene-deleted cell lineages into two groups under each condition (top 50% and bottom 50%) based on their mCherry-CAT fluorescence intensities at the onset of Cp exposure. While we found no significant differences in the *T*_*c*_ = 0 and 3 h conditions, the top 50% group produced 2.4-fold more growth-restored cells than the bottom 50% group in the *T*_*c*_ = 6 h condition (Fig. 3F). In the *T*_*c*_ = 10 h condition, we could barely detect the fluorescence signals from any cells (Fig. 3D and E), and no cells showed growth restoration as mentioned above (Fig. 3F). These results suggest that a low level of residual mCherry-CAT proteins is required at the onset of Cp exposure, but high amounts do not necessarily ensure the growth restoration in all resistance-gene-deleted cells. Moreover, 40% could be considered as the maximum frequencies at which cells can restore growth without the resistance gene.

### Growth restoration accompanies the recovery of stoichiometric balance of ribosomal subunits

The results above demonstrated that the residual mCherry-CAT proteins are important for initiating the transition to growth restoration without the resistance gene. However, the residual proteins cannot directly support the growth in later generations under Cp exposure as their levels are eventually diluted with growth. Indeed, the amounts of mCherry-CAT proteins during growth restoration were below the detection limit (Fig. 2F and H). Therefore, we speculated that the other physiological changes were responsible for growth restoration.

Because Cp targets the 50S ribosomal subunit [17], it is plausible that ribosomal states were modulated in the duration beginning from resistance gene deletion to growth restoration. Hence, we constructed an *E. coli* strain YK0136 that expressed fluorescently-tagged dual ribosomal reporters, RplS-mCherry and RpsB-mVenus (Fig. 4A) [18]. RplS and RpsB are ribosomal proteins in the 50S and 30S subunits, respectively. Their fluorescent reporters were utilized to probe the ribosomal subunit balance in living cells [18]. YK0136 also harbors the *cat* resistance gene (not tagged with *mcherry*) between the upstream and downstream *loxP* sequences on the chromosome, and the PA-Cre recombination system was expressed via the plasmid (Fig. 4A). We confirmed that YK0136 showed MIC values almost equivalent to those of the YK0083 strain (Fig. S3 and S8). Furthermore, 30-min blue-light illuminations provoked *cat*-gene deletion in 28.6% (452/1587) of YK0136 cells, which was comparable to the frequency of *mcherry-cat*-gene deletion in YK0083 (Fig. 3B).

**Figure 4.**
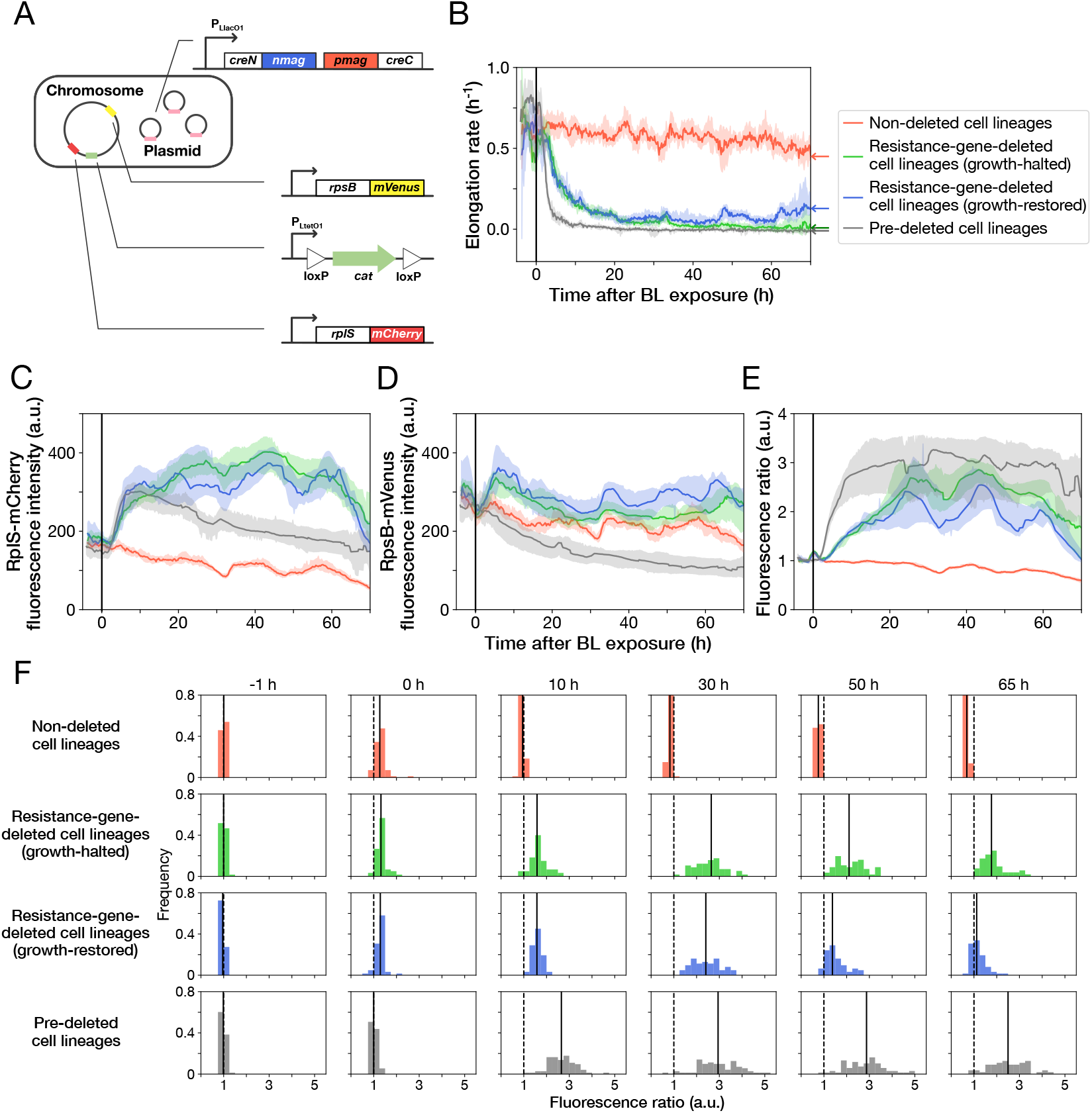
Disruption and restoration of ribosomal proteins’ stoichiometry. (A) Schematic diagram of the ribosome reporter strain, YK0136. Fluorescently-tagged ribosomal protein genes (*rplS* and *rpsB*) are expressed from the native loci on the chromosomes. Additionally, the photo-removable *cat* gene was integrated into the *intC* locus on the chromosome. The PA-Cre fragments were expressed via low-copy plasmids. (B) Transitions of elongation rates. Lines represent the medians of elongation rates at each time point, and shaded areas show the 95% error ranges of the medians estimated by resampling the cell lineages 1,000 times. Red represents non-deleted cell lineages. Green shows growth-halted resistance-gene-deleted cell lineages. Blue represents growth-restored resistance-gene-deleted cell lineages. Gray represents pre-deleted cell lineages. Color correspondence remains the same in the following figures. The end of blue-light illumination was set to 0 h on the horizontal axis. (C-E) Transitions of RplS-mCherry fluorescence intensities (C), RpsB-mVenus fluorescence intensities (D), and the fluorescence ratio of RplS-mCherry/RpsB-mVenus (E). The fluorescence intensities of RplS-mCherry and RpsB-mVenus were normalized by the intensity observed at *t* = 0 h to calculate the ratio in E. (F) Transitions of the RplS-mCherry/RpsB-mVenus fluorescence ratio distributions. The vertical dashed lines represent the position of ratio = 1; the solid lines represent the medians of the distributions.

We conducted time-lapse measurements of YK0136 cells in the mother machine and induced the deletion of the resistance gene via blue-light illumination under continuous exposure to 15 *µ*g/ml of Cp (Movie S4). Since previous experiments showed that the resistance gene deletion decelerates cellular growth significantly, the deletion of the *cat* gene in each cell lineage was evaluated based on the distinctive declines of cellular elongation rates after the blue-light illumination. We validated this criterion by hierarchical time-series clustering applied to the growth of YK0083 cells and mCherry-CAT fluorescence transitions (Fig. S9 and Methods).

As observed previously, 45.9% (158/344) of the *cat*-deleted YK0136 cells gradually restored their growth after initial growth suppression (Fig. 4B, S10 and Movie S4). Furthermore, such growth-restored cell lineages were not observed (0/1138) when YK0138 cells were exposed to 15 *µ*g/ml of Cp; the YK0138 strain was constructed from YK0136 cells by deleting the *cat* gene beforehand in batch culture via blue-light illumination (Fig. 4B and Movie S5).

The transitions of fluorescence intensities reveal that *cat*-gene deletion under Cp exposure increased the expression levels of both RplS-mCherry and RpsB-mVenus (Fig. 4C and D). The relative changes triggered by deletion were more significant for RplS-mCherry than those for RpsB-mVenus (Fig. 4C and D). Consequently, the ribosomal subunit balance probed by the RplS-mCherry/RpsB-mVenus ratio was disrupted, and the ratio increased approximately 3-fold (Fig. 4E and F). The initial disruption kinetics was similar between growth-restored and growth-halted resistance-gene-deleted cell lineages (Fig. 4E). However, the ratio of growth-restored cell lineages gradually returned to the original level, whereas the ratio of growth-halted cell lineages remained disrupted (Fig. 4E and F). These results suggest that the regain of the ribosomal subunit balance under Cp exposure is required for growth restoration.

Analysis of the correlations between the pre-illumination phenotypic traits and post-illumination cellular fates reveals that elongation rates, fluorescence intensities of RplS-mCherry and RpsB-mVenus, and the fluorescence ratio observed before blue-light illumination did not strongly affect the likelihood of gene deletion and the determination of growth-halting and growth-restoration fates (Fig. S11).

When non-resistant YK0138 cells were exposed to 15 *µ*g/ml of Cp, the expression levels of RplS-mCherry increased initially following kinetics similar to those of the YK0136 cells observed after *cat*-gene deletion (Fig. 4C). In contrast, the expression levels of RpsB-mVenus decreased in response to Cp exposure (Fig. 4D), as previously reported for a wildtype *E. coli* strain exposed to Cp [19]. The initial decrease of RpsB-mVenus expression led to a more rapid disruption of the RplS-mCherry/RpsB-mVenus ratio in the susceptible YK0138 cells (Fig. 4E and F). Consistently, the decline in the elongation rates of the YK0138 cells was faster than those of the YK0136 cells (Fig. 4B).

We further examined the relationship between the RplS-mCherry/RpsB-mVenus ratio and elongation rates and observed a negative correlation (Spearman *ρ* = -0.58, 95% confidence interval [-0.62,-0.53]; Fig. 5). The ratio was maintained within a narrow range (0.84-1.18, 95% interval) before blue-light illumination (Fig. 4F and 5). The disruption of ribosomal subunit balance exceeding this range was linked to growth suppression (Fig. 5). These results suggest that the ribosomal subunit balance is essential for growth under Cp exposure and that cells must restore the balance to recover growth in the absence of the resistance gene.

**Figure 5.**
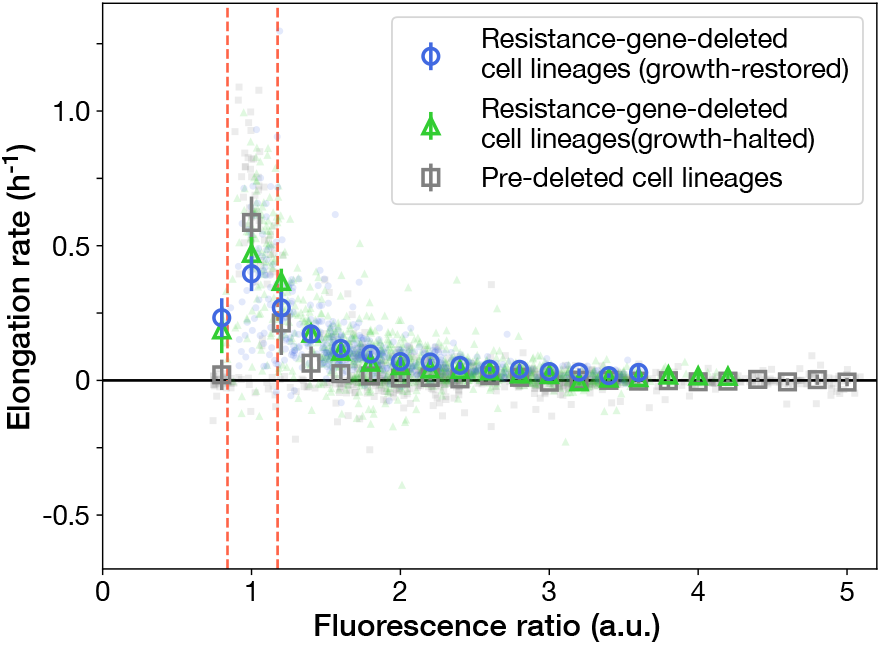
Scatter plot of fluorescence ratio and elongation rate under Cp exposure. Small points represent the relations between the fluorescence ratio and elongation rates averaged over the two-hour periods at the different time points on the single-cell lineages. The larger open point represents the mean elongation rate in each bin of fluorescence ratio (bin width, 0.2 a.u.). Error bars represent the 1.96 × standard error ranges. Blue represents growth-restored resistance-gene-deleted cell lineages. Green represents growth-halted resistance-gene-deleted cell lineages. Gray represents pre-deleted cell lineages.Red dashed lines indicate the mean ± 1.96× standard error range of fluorescence ratio of non-deleted cell lineages before blue-light illumination.

## Discussion

Uncovering the genotype-phenotype correspondences in various biological contexts is the groundwork for genetics. However, the correspondences are usually investigated based on terminal phenotypes, which are often manifested after significant lags following genotypic changes. Consequently, our understanding of the dependence of terminal phenotypes on historical conditions and the heterogeneity of phenotypic consequences among individual cells remains limited. In this study, we demonstrated that *E. coli* cells could adapt even to a lethal genetic modification, i.e., the deletion of *cat* resistance gene under exposure to chloramphenicol (Fig. 2). Importantly, such adaptation was not observed when an identical genetic modification was introduced long before Cp exposure (Fig. 3). Therefore, whether cells could gain physiological resistance in the absence of a resistance gene depends on the timing of resistance gene deletion and environmental histories that the cells experienced.

The analyses revealed that the *cat* gene deletion under Cp exposure disrupted the stoichiometric balance of ribosomal proteins (RplS and RspB) and that the balance was gradually recovered alongside growth restoration (Fig. 4). The elongation rate and the stoichiometric balance of ribosomal proteins were strongly correlated (Fig. 5). Furthermore, the balance was tightly regulated within a narrow range in the fast-growing non-deleted cells (Fig. 4F). Therefore, the recovery of ribosomal stoichiometric balance might be essential for restoring growth against the deletion of the resistance gene.

It still remains elusive how individual cells recovered the ribosomal stoichiometric balance under continuous Cp exposure. A plausible scenario might be that the multi-layered and interlinked feedback regulations on ribosomal components entailed stoichiometry readjustments [20–24]. Many ribosomal proteins bind to their own mRNAs, in addition to the target ribosomal RNAs [21–24]. These regulatory fractions of ribosomal proteins bind to their mRNAs and inhibit their own translation and that of other genes in the same operons. The competition among rRNA and mRNA for these regulatory ribosomal protein components is considered to regulate the stoichiometry of ribosomal components [21–24]. Therefore, it is plausible that these innate homeostatic mechanisms of ribosomes might have been exploited by cells to restore growth. It is also important to note that the operons for ribosomal proteins comprise genes encoding DNA polymerase subunits, initiation factors, and protein transmembrane transporters [22, 23]. Hence, the feedback regulations in ribosomes could mediate changes in the global expression profiles, which might have helped the cells find growth-permissive states in the presence of Cp.

The necessity for a small amount of residual resistant proteins at the onset of Cp exposure (Fig. 3) might suggest that experiencing such barely-tolerable intracellular states is crucial for gaining physiological resistance. The importance of near-critical states for survival might be related to the phenomenon that pre-treatment with a sublethal concentration of antibiotics increases the fraction of persister cells in response to the subsequent treatments with lethal antibiotic concentrations [25, 26]. Whether analogous mechanisms are involved in adaptation to both internal and external lethal perturbations is worth examining.

Though experimental evidence of history-dependent physiological adaptation to lethal genetic perturbations is still limited, the experimental strategy combining optogenetic recombination and single-cell lineage tracking is applicable for the modification of other genes or other cell types, including mammalian cells. Such approaches might offer new insights into the emergence of drug-resistant bacterial and cancer cells. Gaining a more comprehensive picture of resistance evolution might help us design new drug treatment strategies to counter or control the emergence and spread of resistance.

## Acknowledgments

We thank Kaito Kikuchi for the help with plasmid construction in the initial stage of this project; Stanislas Leibler, Chikara Furusawa, Kunihiko Kaneko, Tetsuya J. Kobayashi, and the members of the Wakamoto Lab for discussion. This work was supported by JST CREST Grant Number JPMJCR1927 (Y.W.) and JPMJCR1653 (M.S.); JST ERATO Grant Number JPMJER1902 (Y.W.); Japan Society for the Promotion of Science KAKENHI Grant Number 17H06389 and 19H03216 (Y.W.); Project Grant from Kanagawa Institute of Industrial Science and Technology (KISTEC) (M. S.); and Grant-in-Aid for JSPS Fellows Grant Number JP19J22506 (Y.K.).

## Materials and Methods

### Bacterial strains, plasmids construction, and culture conditions

*E. coli* strains, plasmids, and primers are listed in Table S2, S3, and S4, respectively. For strain constructions, cells were grown in Luria Bertani (LB) broth (Difco) at 37°C unless stated otherwise. Plasmids were constructed using DNA ligase (T4 DNA Ligase, Takara) or by ExoIII cloning [27] and introduced into the *E. coli* strain JM109 for cloning and stocking. The *cat* or *mcherry-cat* gene was introduced into the *intC* locus of F3 (W3110Δ*fliC* ::FRTΔ*fimA*::FRTΔ*flu*::FRT) genome by *λ*-Red recombination (YK0080, YK0134) [28]. Fluorescently-tagged ribosome reporter genes were introduced into the *E. coli* strain BW25113 (MUS3, MUS13). These ribosome reporter genes were transferred into the F3 strain by P1 transduction (MUS5, MUS6). The colonies with intended genome integration were selected on LB plates containing kanamycin (Km, Wako). The Km resistant gene was removed by flp-FRT recombination using pCP20 plasmid [28].

When constructing the resistance-gene-deleted strains (YK0085 and YK0138), YK0083 and YK0136 cells were pre-cultivated overnight in LB broth containing 50 *µ*g/mL of ampicillin (Amp, Wako). 10 *µ*L of overnight cultures were inoculated into 2 mL of the M9 medium containing 50 *µ*g/mL of Amp and 0.1 mM isopropyl-*β*-D-thiogalactopyranoside (IPTG, Wako) in test tubes. The cell cultures were cultivated at 37°C with shaking. After a 3 h cultivation period for inducing the expression of PA-Cre system by IPTG, the batch cultures were illuminated by LED blue light (CCS) for 24 h. The light intensity was adjusted to 6.8 mW, which is equivalent to the intensity in the microscopy experiments. The cell cultures were streaked on LB agar containing 100 *µ*g/mL of Amp. The plates were incubated at 37°C overnight. After overnight incubation, several colonies were selected. The deletion of the intended gene in these colonies was examined via PCR. The colonies with the intended gene deletion were cultured in LB medium containing 50 *µ*g/mL of Amp at 37°C. These cell cultures were stored at −80°C as glycerol stocks.

The genomes of the constructed strains and plasmids were purified using the Promega Genomic DNA Purification kit and the Promega Plasmid Miniprep System, respectively, and the DNA sequence of the modified locus was amplified by Prime STAR (Takara). The sequence was examined by Sanger sequencing using a commercial service (FASMAC).

In microscopy experiments, we used M9 minimal medium, which consisted of M9 salt (Difco), 2 mM MgSO_4_ (Wako), 0.1 mM CaCl_2_ (Wako), 0.2% (w/v) glucose (Wako) and 0.2%(v/v) MEM amino acid (50x) solution (Sigma). When necessary, antibiotics and chemicals were added at the following final concentrations unless otherwise noted: Amp 50 *µ*g/mL, Km 20 *µ*g/mL, chloramphenicol (Cp, Wako) 15 *µ*g/mL, and IPTG 0.1 mM.

### Calculation of the fraction of resistance-gene-deleted cells in batch culture

YK0083 cells were pre-cultivated in LB broth containing 50 *µ*g/mL of Amp overnight. 100 *µ*L of the overnight culture was spun at 21,500 × *g* for 1 minute by a centrifuge (CT15E, himac, Hitachi) and resuspended in 1 mL of the M9 minimal medium. This culture was inoculated in 2 mL of the M9 medium containing 50 *µ*g/mL of Amp and 0.1 mM IPTG (except for the “no IPTG” condition) in test tubes covered with aluminum foil for light protection. The starting OD_600_ was adjusted to 0.001, and the cells were cultivated at 37°C with shaking. After a 3 h cultivation period for inducing the expression of the PA-Cre system by IPTG, the aluminum foil cover was removed, and the test tubes were exposed to blue light. The light intensity was adjusted to 6.8 mW. The illumination length was determined according to the experimental conditions (Fig. S2B). For the conditions with an illumination duration of less than 6 h, the test tubes were again covered with aluminum foil and incubated with shaking so that the total cultivation duration from the start of blue-light illumination was identical (6 h) across all conditions. Under the “no IPTG” condition, the test tubes were covered with the aluminum foil throughout the incubation period and were not exposed to blue light.

For calculating the fractions of resistance-gene-deleted cells, the cultures exposed to blue-light were diluted to OD_600_ = 1.0 × 10^−6^, and 150 *µ*L of the diluted cultures was spread on LB agar containing 100 *µ*g/mL of Amp. The plates were incubated at 37°C for 18 h. After incubation, the number of colonies was counted under ambient light or excitation light for examining mCherry fluorescence using stereomicroscope (stereomicroscope: Olympus SZ61; LED source: NIGHTSEA SFA-GR). The fraction of resistance-gene-deleted cells was calculated as the number of non-fluorescent colonies divided by the number of total colonies (Fig. S2B). To validate this classification, we conducted colony PCR and checked whether the designed deletion was introduced (Fig. S2C).

### Sample preparation and experimental procedures for MIC tests

Cells stored as a glycerol stock were inoculated in the LB broth (Fig. S3, and S8) or the M9 medium (Fig. S6C) containing 50 *µ*g/mL of Amp. The cells were cultured at 37°C with shaking overnight. 100 *µ*L of overnight culture was centrifuged at 21,500×*g* for 3 min. The supernatant was discarded, and the cells were resuspended in 1.0 mL of the M9 medium. This culture was diluted to the cell density corresponding to OD_600_ = 0.01 in the M9 medium containing 50 *µ*g/mL of Amp and incubated with shaking for 3-4 h at 37°C. The cell culture was again diluted to OD_600_ = 0.001 in fresh M9 media containing different concentrations of Cp (Fig. S3, S6C and S8). After 23 h of incubation, the OD_600_ of these cultures was measured with a spectrometer (UV-1800, Shimadzu) in Fig. S3 and S8 or with a multi-mode microplate reader (FilterMax F5, Molecular Devices) in Fig. S6C.

### Microfabrication of Mother Machine microfluidic device

We created two chromium photomasks, one for the main trench and the other for the observation channels, by laser drawing (DDB-201-TW, Neoark) on mask blanks (CBL4006Du-AZP, Clean Surface Technology). The photoresist on mask blanks was developed in NMD-3 (Tokyo Ohka Kogyo), and the uncovered parts of the chromium layer were removed by MPM-E30 (DNP Fine Chemicals). The remaining photoresist was removed by acetone. The photomasks were rinsed in MilliQ water and air-dried.

We created the mold for the mother machine on a silicon wafer (ID447, *ϕ*=76.2 mm, University Wafer). First, we spin-coated SU8-2 (MicroChem) on the wafer with the target height of 1.2 *µ*m. The SU8-coated wafer was baked at 65°C for 1 min and thereafter at 95°C for 3 min. The SU8-layer was exposed to UV light thrice (each exposure, 22.4 mW/cm^2^, 1.7 sec) using a mask aligner (MA-20, Mikasa). The photomask for the observation channels was used in this step. The wafer was post-baked at 65°C for 1 min and 95°C for 3 min after the exposure and developed with the SU8 developer. The wafer was rinsed with isopropanol (Wako) and air-dried.

Next, we spin-coated SU8-3025 (MicroChem) with the target height of 20 *µ*m. The pre-bake was performed at 65°C for 3 min and thereafter at 95°C for 7 min. The SU-8 layer was again exposed to UV light (22.4 mW/cm^2^, 30 sec) with the photomask for the main trench. The wafer was post-baked at 65°C for 3 min and 95°C for 10 min and developed with the SU8 developer. The wafer was rinsed with isopropanol and air-dried.

The Part A and Part B of polydimethylsiloxane (PDMS) resin (Sylgard 184 Silicone Elastomer Kit, Dow Corning) were mixed at the ratio of 10:1 and poured onto the SU-8 mold placed in a tray made of aluminum foil. After removing the air bubbles under decreased pressure, the PDMS was cured at 65°C for 1 h. We cut out a PDMS block containing the channel structures and punched holes at both ends of the main trench. The PDMS block was washed with isopropanol and heated at 65°C for 30 min.

We washed the coverslips (thickness: 0.13-0.17 mm, 24 × 60 mm, Matsunami) by sonication with 10-fold-diluted Contaminon solution (Contaminon ® LS-II, Wako) for 30 min, with 99.5% ethanol (Wako) for 15 min, and with 0.8 M NaOH solution (10-fold diluted 8 M NaOH (Wako)) for 30 min. The coverslips were rinsed with milliQ water by sonication after each washing step. The coverslips were dried at 140°C for 1 h.

We exposed the PDMS block and the coverslips to oxygen plasma using a compact etcher (FA-1, SAMCO) and bonded them together at 65°C for 5 min. After bonding, we inserted the silicone tubes (inner diameter: 1 mm, outer diameter: 2 mm, Tigers Polymer Corporation) into the holes and smeared a small amount of pre-cured PDMS. We cured the PDMS at 65°C overnight to fix the tubes into the holes tightly.

### Sample preparation and experimental procedures for single-cell time-lapse experiments

Cells stored as a glycerol stock were inoculated in the LB broth containing 50 *µ*g/mL of Amp and cultured at 37°C overnight with shaking (200 rpm). 200 *µ*L of overnight culture was centrifuged at 21,500 × *g* for 3 min. The supernatant was discarded, and the cells were resuspended in 1.5 mL of the M9 medium. This culture was diluted to OD_600_ = 0.01 in the M9 medium containing 50 *µ*g/mL of Amp, and incubated with shaking for 5-6 h at 37°C. The cell culture was again spun at 2,350 × *g* for 5 min using a centrifuge (CT6E, himac, Hitachi), and the cells were resuspended in 200 *µ*L of the M9 medium.

Before loading the cells, the growth channels and the main trench in the mother machine device were washed by flowing 0.5 mL of 99.5% ethanol, M9 medium, and 1% (w/v) bovine serum albumin (BSA, Wako) solution sequentially using a syringe. After washing, we introduced the cell suspension into the device and let the cells enter the growth channels by placing the device at 37°C for 1-2 h without flow in a dark room. After confirming that the cells were trapped in many growth channels, we started flowing the M9 medium containing 0.1 mM IPTG and 0.1% (w/v) BSA at 2 mL/h. For experiments with continuous Cp exposure, we also added 15 *µ*g/mL of Cp to the flowing medium from the beginning. We initiated time-lapse image acquisitions after overnight cultivation in the mother machine. We illuminated blue light for 30 min at pre-determined timings in all single-cell time-lapse experiments. The medium flow rate was maintained at 2 mL/h and increased to 5 mL/h for 30 min once or twice a day to prevent the formation of cell aggregates in the main trench.

### Time-lapse image acquisitions and analysis

Time-lapse image acquisitions were performed with ECLIPSE Ti fluorescent microscope (Nikon) equipped with 100 × oil immersion objective lens (Plan Apo *λ*, NA 1.45, Nikon), digital CMOS camera (ORCA-frash, Hamamatsu Photonics), and LED light source (DC2100, Thorlabs) for fluorescence excitation. For acquiring phase-contrast images, the transmitted light was illuminated for 20 msec during *mcherry-cat* gene deletion experiments and for 50 msec during experiments performed with ribosome reporter strains through neutral density and red filters to avoid unintended activation of PA-Cre. Fluorescence images were acquired with appropriate filter cubes (YFP HQ (Nikon) for mVenus and Texas Red (Nikon) for mCherry). In the *mcherry-cat* gene deletion experiments, mCherry fluorescence was obtained every 2 min with excitation of 500 msec. The mCherry and mVenus fluorescence images were acquired every 10 min with the excitation of 100 msec in the experiments with ribosome reporter strains. IPTG was removed from the flowing media after blue-light illumination to avoid unintended gene deletion.

Blue-light illumination on the microscope for gene deletion experiments was regulated by an independent controller. We used a tape LED (7.2 mW, wave length:464 ∼ 474 nm, 60 LED/1 m, LED PARADISE) as the blue-light source and surrounded the mother machine device on the microscope stage with this tape LED. The LED light was switched on and off with a timer (REVEX).

Image analysis was performed using ImageJ Fiji (http://fiji.sc/). Image registration was performed by HyperStackReg described in MoMA Macro [29]. Cell segmentation was semi-automated and retouched manually using iPad Pro Sidecar. Semi-automated segmentation and tracking Macro reported previously [30] were used in this study. The data from image analysis were further analyzed using Python 3 (https://www.python.org/) with some general packages, including NumPy, SciPy, pandas, Matplotlib, seaborn, FastDTW, and JupyterLab.

The transitions of mCherry-CAT fluorescence intensities and elongation rates in single-cell lineages were classified using hierarchical clustering in Fig. S9. The full-length single-cell lineage data were used in this analysis. The similarity of each pair of time series was measured by dynamic time warping [31]. Hierarchical clustering is agglomerative; the averaged dynamic time warping was used when similarity to another time series or cluster was evaluated.

### Cell sampling from the mother machine and whole-genome sequencing

After time-lapse observations performed with the mother machine, we switched the flowing media to the M9 medium containing 0.1% (w/v) BSA and no Cp. We recovered the growth of growth-restored resistance-gene-deleted cell lineages by culturing them without Cp for more than 6 h. Subsequently, we collected the media flowing out from the mother machine in a 1.5-mL tube. The OD_600_ of the collected cell suspension was measured with a spectrometer (UV-1800, Shimadzu). The cell suspension was diluted serially with the M9 medium to obtain a cell density corresponding to OD_600_ = 3.3 × 10^−9^. 200 *µ*L of diluted cell suspension was taken in each well of 96-well plates. The number of cells in the 200 *µ*L of cell suspension *k* is expected to follow a Poisson distribution *P* (*k*) = *λ*^*k*^*e*^−*λ*^*/k*! with the mean *λ* = 0.54. Therefore, *P* (0) = 0.58, *P* (1) = 0.31, *P* (2) = 0.085, and *P* (*k≥* 3) = 0.018. The 96-well plates were incubated with shaking at 37°C for two nights. We performed PCR on samples with high turbidity in wells to confirm the presence or absence of the *cat* gene. We selected both *cat*-positive and *cat*-negative cellular populations from the 96-well plates and cultured them in test tubes overnight at 37°C. We stored these samples at −80°C for further analyses.

For the measurement of mCherry-CAT fluorescence intensities of the selected samples (Fig. S6), cells stored as a glycerol stock were inoculated in the M9 medium containing 50 *µ*g/mL of Amp and cultured at 37°C with shaking (200 rpm) overnight. 10 *µ*L of overnight culture was inoculated in the M9 medium containing 50 *µ*g/mL of Amp and incubated with shaking for 4 h at 37°C. 0.3 *µ*L of cell culture was placed on an agar pad prepared using the M9 medium and 1.5% (w/v) agar (Wako) and covered with a coverslip. Image acquisitions of the samples were performed with ECLIPSE Ti fluorescent microscope (Nikon) equipped with 100 × oil immersion objective lens (Plan Apo *λ*, NA 1.45, Nikon), digital CCD camera (ORCA-R2, Hamamatsu Photonics), and LED light source (DC2100, Thorlabs) for fluorescence excitation. For acquiring phase-contrast images, the transmitted light was illuminated for 50 msec through neutral density. The mCherry fluorescence images were acquired with Texas Red filter cubes and an exposure time of 500 msec.

For the measurement of whole-genome sequencing of the selected samples, cells stored as a glycerol stock were inoculated in 5 mL of the M9 medium and cultured at 37°C with shaking (200 rpm) overnight. The OD_600_ of overnight culture was measured with a spectrometer. Rifanpicin (Wako) was added to the overnight culture to the final concentration of 300 *µ*g/mL. After 3 h of incubation, the genomes of these samples were extracted with DNeasy Blood and Tissue kit (QIAGEN). TruSeq DNA PCR Free kit (Illumina) was used for the whole-genome sequencing library preparation. These libraries were sequenced using NovaSeq (Illumina). Library preparation and sequencing were outsourced to Macrogen Japan (Tokyo, Japan). The sequence data were analyzed with breseq [32].

## Supplemental Information

### Supplemental Movies

**Movie S1. Deletion of *mcherry-cat* gene by blue-light illumination under drug-free conditions**. YK0083 cells were cultured in the mother machine flowing the M9 minimal medium and exposed to blue light from *t* = −30 min to 0 min (marked with “+ BL”). The merged images of phase contrast (grayscale) and mCherry-CAT fluorescence (red) channels are shown. The fluorescence of mCherry-CAT was gradually lost after blue-light illumination in this cell lineage. Scale bar, 5 *µ*m.

**Movie S2. Growth restoration under Cp exposure after resistance gene deletion**. YK0083 cells were cultured in the mother machine flowing the M9 minimal medium containing 15 *µ*g/mL of Cp. Blue light was illuminated from *t* = −30 min to 0 min (marked with “+ BL”). The merged images of phase contrast (grayscale) and mCherry-CAT fluorescence (red) channels are shown. The illumination caused the loss of fluorescence signals, i.e., the deletion of the *mcherry-cat* gene, in this cell lineage. Nevertheless, the cell gradually restored growth and division under continuous Cp exposure. Scale bar, 5 *µ*m.

**Movie S3. No growth recovery of YK0085 strain under Cp exposure**. YK0085 cells were cultured in the mother machine flowing the M9 minimal medium and exposed to 15 *µ*g/mL of Cp at *t* = 0 min (marked with “+ Cp”). The Cp exposure caused growth arrest, and, unlike the YK0083 cells, the cell did not restore growth. We found no growth-restored cell lineages in this experimental condition. Scale bar, 5 *µ*m.

**Movie S4. Recovery of the balance of ribosomal proteins alongside growth restoration**. YK0136 cells were cultured in the mother machine flowing the M9 minimal medium containing 15 *µ*g/mL of Cp. Blue light was illuminated from *t* = −30 min to 0 min (marked with “+ BL”). The merged images of phase contrast (grayscale), RplS-mCherry fluorescence (red), and RpsB-mVenus fluorescence (green) channels are shown. The growth of the cell markedly slowed down after blue-light illumination, indicating the loss of the *cat* gene. The cell became red in response to blue-light illumination, indicating that the relative expression level of RplS-mCherry became higher than that of RpsB-mVenus. However, the balance was restored alongside growth restoration. Scale bar, 5 *µ*m.

**Movie S5. No recovery of the balance of ribosomal proteins in the YK0138 strain**. YK0138 cells were cultured in the mother machine flowing the M9 minimal medium and exposed to 15 *µ*g/mL of Cp at *t* = 0 min (marked with “+ Cp”). The Cp exposure caused growth arrest and disruption of ribosomal proteins’ expression balance. No restoration of growth and ribosomal balance was observed. Scale bar, 5 *µ*m.

### Supplemental Tables

**Table S1.**
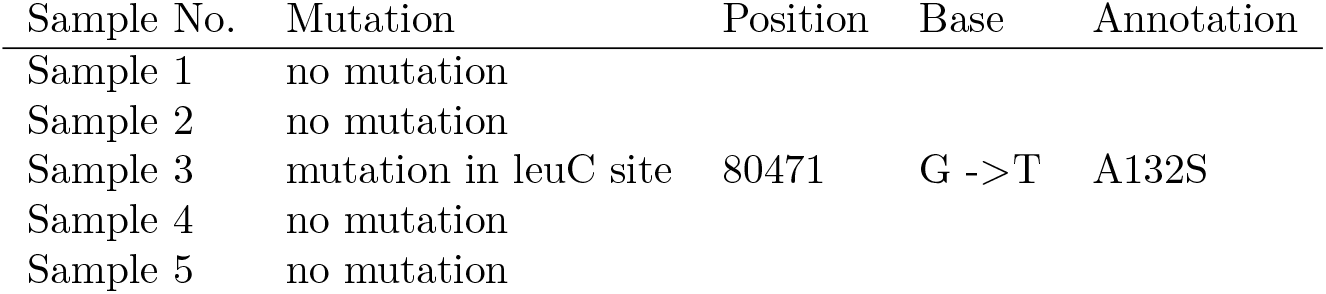
Mutations detected by whole-genome sequencing. We obtained isolated cellular populations derived from single or few cells by limiting dilution of the culture media flowing out from the mother machine. We detected only one point mutation in Sample 3.

| Sample No. | Mutation | Position | Base | Annotation |
| --- | --- | --- | --- | --- |
| Sample 1 | no mutation |  |  |  |
| Sample 2 | no mutation |  |  |  |
| Sample 3 | mutation in leuC site | 80471 | G ->T | A132S |
| Sample 4 | no mutation |  |  |  |
| Sample 5 | no mutation |  |  |  |

**Table S2.** Strain list used in this study.

mCherry-CAT deletion
| Name | Genotype | Source |
| --- | --- | --- |
| F3 | W3110 $\Delta fliC::FRT\Delta fimA::FRT\Delta flu::FRT$ | [30] |
| YK0080 | F3 *intC::P<sub>LtetO1</sub>-loxP-RBS4-mcherry-cat-loxP-FRT* | This study |
| YK0083 | F3 *intC::P<sub>LtetO1</sub>-loxP-RBS4-mcherry-cat-loxP-FRT/pYK022* | This study |
| YK0085 | F3 *intC::P<sub>LtetO1</sub>-FRT/pYK022* | This study |

Ribosome reporter
| Name | Genotype | Source |
| --- | --- | --- |
| MUS3 | BW25113 *rplS-mcherry-FRT-kan-FRT* | This study |
| MUS13 | BW25113 *rpsB-mvenus-FRT-kan-FRT* | This study |
| MUS5 | F3 *rplS-mcherry-FRT* | This study |
| MUS6 | F3 *rplS-mcherry-FRT rpsB-mvenus-FRT* | This study |
| YK0134 | MUS6 *intC::P<sub>LtetO1</sub>-loxP-RBS4-cat-loxP-FRT* | This study |
| YK0136 | MUS6 *intC::P<sub>LtetO1</sub>-loxP-RBS4-cat-loxP-FRT/pYK022* | This study |
| YK0138 | MUS6 *intC::P<sub>LtetO1</sub>-loxP-FRT/pYK022* | This study |

**Table S3.** Plasmid list used in this study.

| Name | Backbone | Gene | Source |
| --- | --- | --- | --- |
| pKD46 |  |  | CGSC |
| pCP20 |  |  | CGSC |
|  | pcDNA3.1 | pCMV-lox2272-loxp-inversed *mvenus-lox2272-loxp* | Sato lab |
|  | pcDNA3.1 | pCMV-rfp-*pmag-creC* | Sato lab |
|  | pcDNA3.1 | pCMV-rfp-*nmag-creN* | Sato lab |
| pTmCherryK3 | pMW118 | P<sub>LtetO1</sub>-RBS3-*mcherry*-FLP-*kan*-FLP | Wakamoto lab |
| pLVK4 | pMW118 | P<sub>LlacO1</sub>-RBS4-*mvenus*-FLP-*kan*-FLP | Wakamoto lab |
| pTVCK4 | pMW118 | P<sub>LtetO1</sub>-RBS4-*mvenus-cat*-FLP-*kan*-FLP | Wakamoto lab |
| pKK1 | pMW118 | P<sub>LlacO1</sub>-rfp-*pmag-creC*-FLP-*kan*-FLP | This study |
| pKK2 | pMW118 | P<sub>LlacO1</sub>-rfp-*nmag-creN*-FLP-*kan*-FLP | This study |
| pYK001 | pMW118 | P<sub>LlacO1</sub>-RBS3-*pmag-creC*-FLP-*kan*-FLP | This study |
| pYK002 | pMW118 | P<sub>LlacO1</sub>-RBS3-*nmag-creN*-FLP-*kan*-FLP | This study |
| pYK006 | pMW118 | P<sub>LtetO1</sub>-RBS3-lox2272-*mvenus-loxp*-FLP-*kan*-FLP | This study |
| pYK007 | pMW118 | P<sub>LtetO1</sub>-RBS3-lox2272-*mcherry-loxp*-FLP-*kan*-FLP | This study |
| pYK008 | pMW118 | P<sub>LtetO1</sub>-RBS3-lox2272-*mcherry-loxp*-FLP-*kan*-FLP | This study |
| pYK009 | pMW118 | P<sub>LtetO1</sub>-RBS3-lox2272-*mcherry-loxp*-FLP-*kan*-FLP | This study |
| pYK010 | pMW118 | P<sub>LtetO1</sub>-RBS4-lox2272-*mcherry-loxp*-FLP-*kan*-FLP | This study |
| pYK011 | pMW118 | P<sub>LtetO1</sub>-lox2272-RBS4-*mcherry-loxp*-FLP-*kan*-FLP | This study |
| pYK016 | pMW118 | P<sub>LlacO1</sub>-RBS3-*pmag-creC*-FLP-*kan*-FLP | This study |
| pYK017 | pMW118 | P<sub>LlacO1</sub>-RBS3-*creN-nmag*-FLP-*kan*-FLP | This study |
| pYK018 | pMW118 | P<sub>LlacO1</sub>-RBS3-*creN-nmag-pmag-creC*-FLP-*kan*-FLP | This study |
| pYK022 | pMW118 | P<sub>LlacO1</sub>-RBS3-*creN-nmag-pmag-creC*-FLP-*kan*-FLP | This study |
| pYK023 | pMW118 | P<sub>LtetO1</sub>-loxp-RBS4-*mcherry-loxp*-FLP-*kan*-FLP | This study |
| pYK028 | pMW118 | P<sub>LtetO1</sub>-loxp-RBS4-*mcherry-loxp*-FLP-*kan*-FLP | This study |
| pYK029 | pMW118 | P<sub>LtetO1</sub>-loxp-RBS4-*mcherry-cat-loxp*-FLP-*kan*-FLP | This study |
| pYK035 | pMW118 | P<sub>LtetO1</sub>-loxp-RBS4-*cat-loxp*-FLP-*kan*-FLP | This study |
| pMU1 | pMW118 | P<sub>LlacO1</sub>-RBS4-*mcherry*-FLP-*kan*-FLP | This study |
| pMU2 | pMW118 | P<sub>LlacO1</sub>-RBS4-*mvenus*-FLP-*kan*-FLP | This study |

**Table S4.**
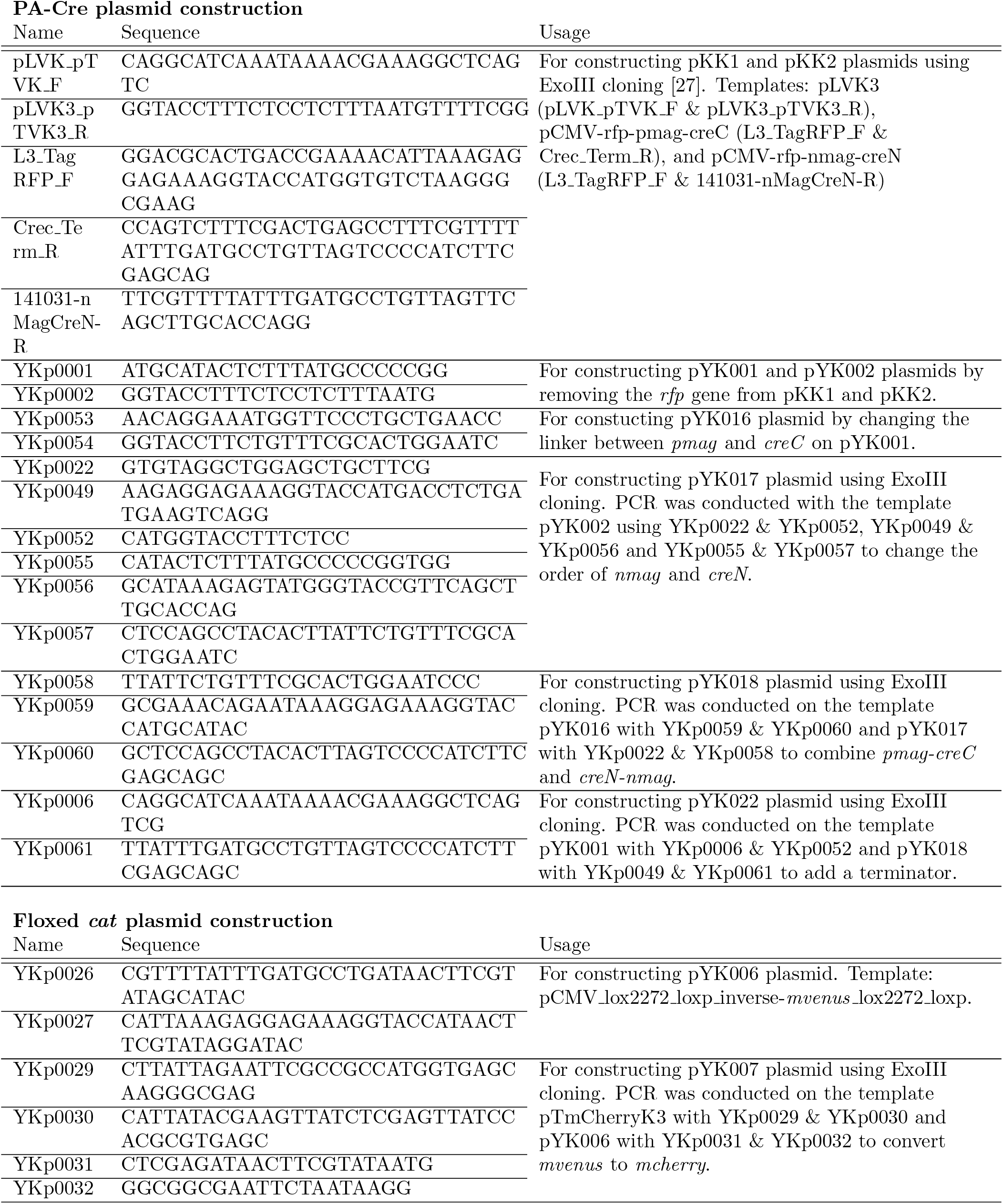
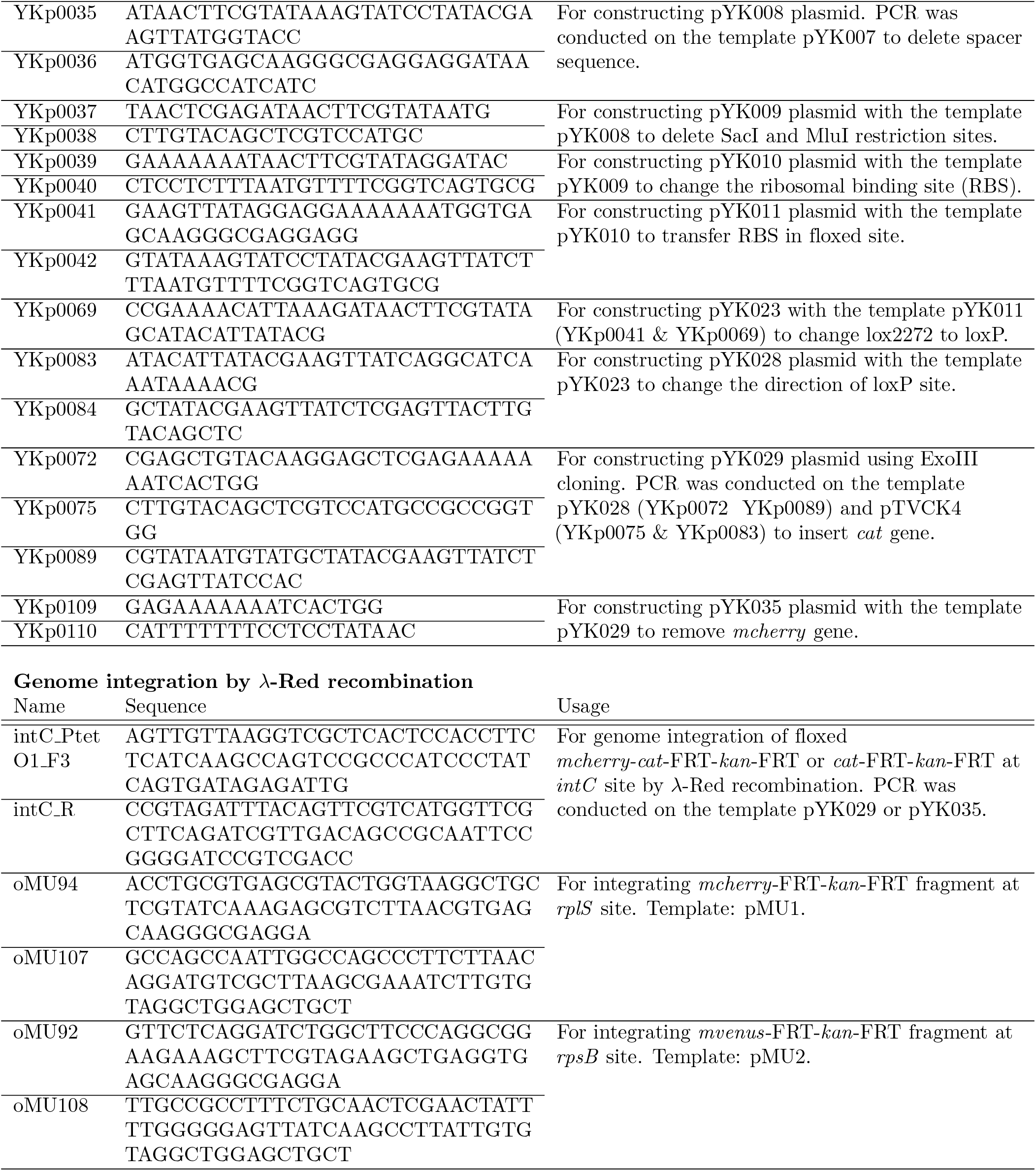
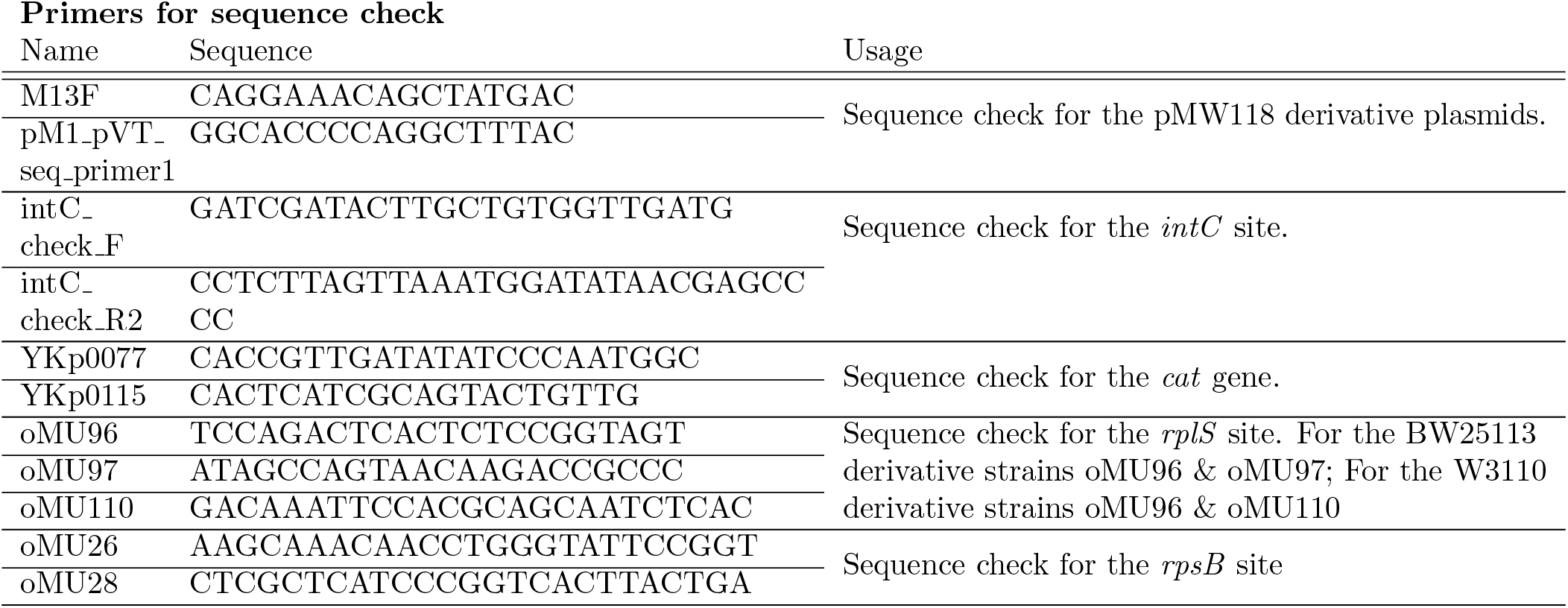
Primer list used in this study.

### Supplemental Figures

**Figure S1.**
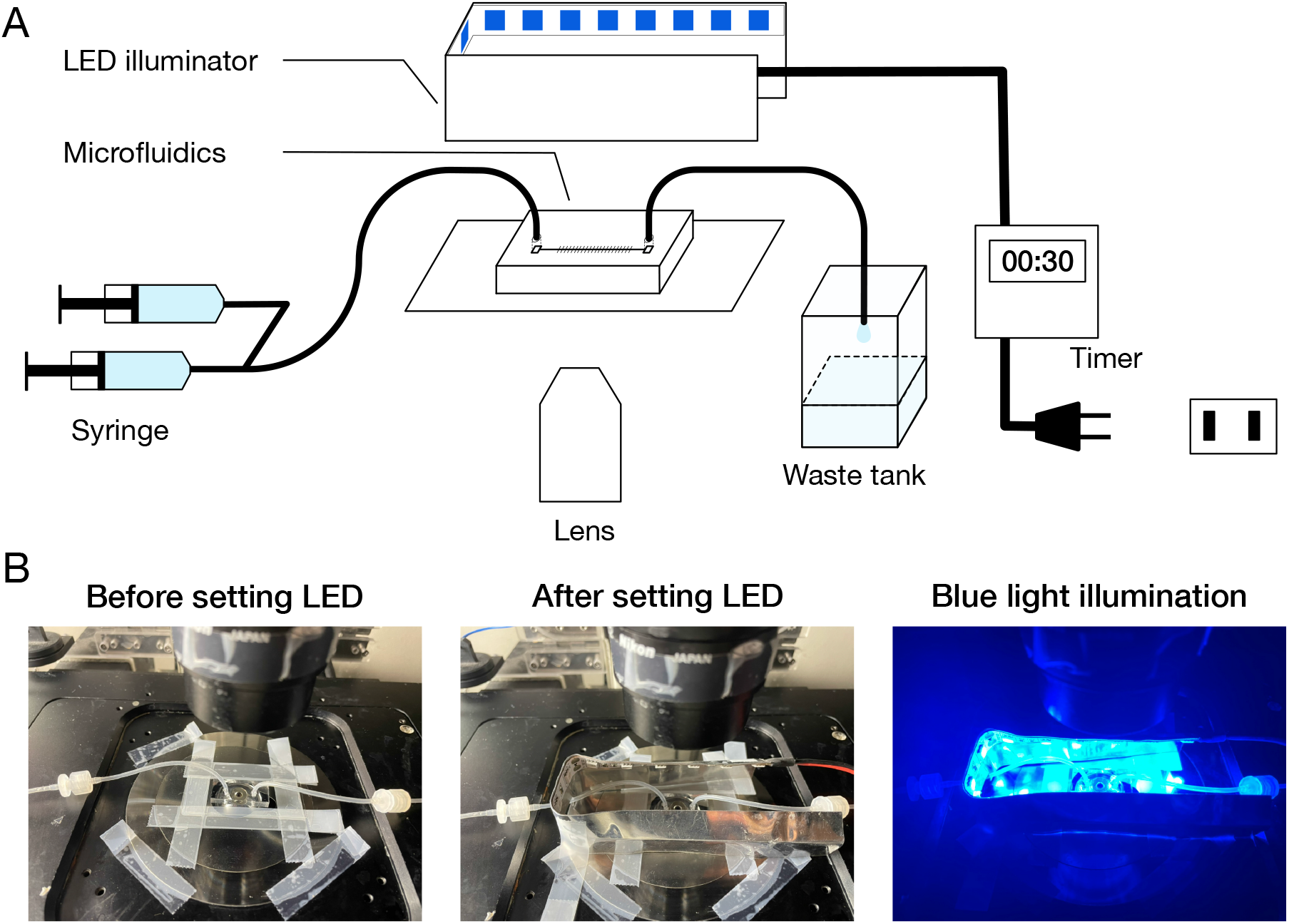
On-stage blue-light LED illuminator. (A) Schematic diagram of microscope setup. A custom blue-light LED illuminator was placed on the microscope stage to illuminate the microfluidic device. The timing and duration of illumination were controlled by an external timer. (B) Photo images of the microfluidic device and LED illuminator mounted on the microscope stage.

**Figure S2.**
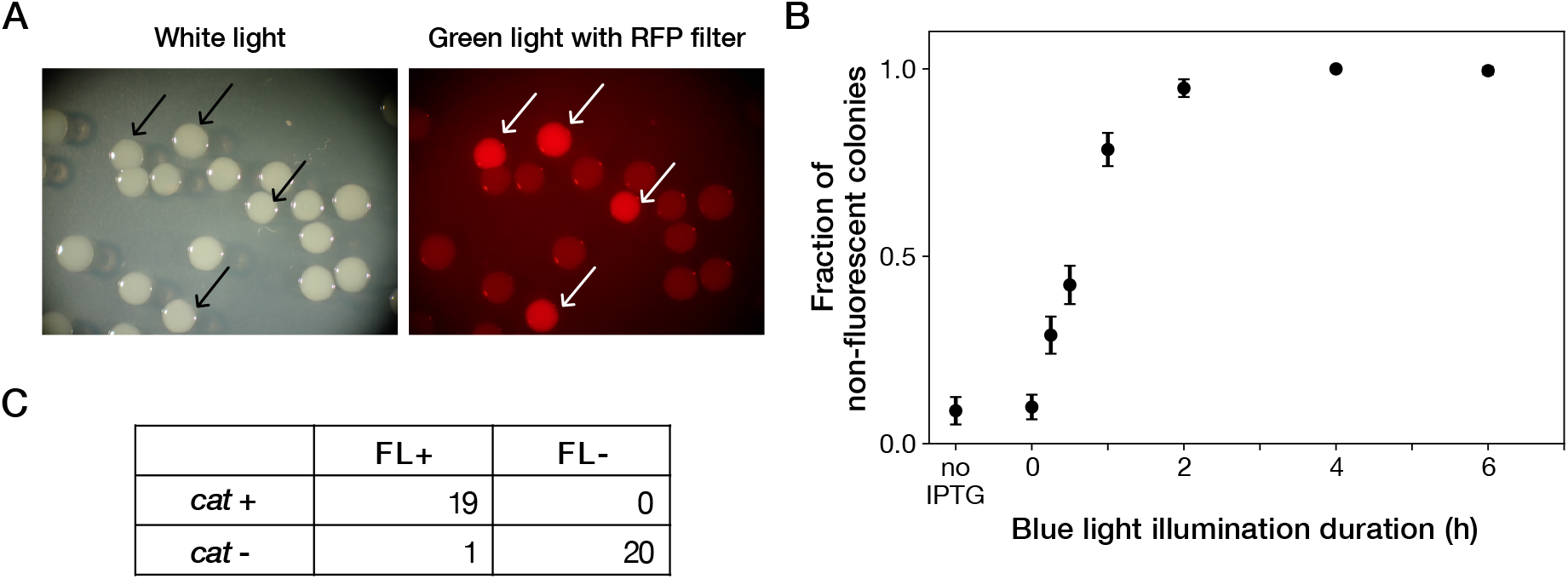
Resistance gene deletion by blue-light illumination in batch cultures. (A) Colonies of YK0083 cells inoculated after blue-light illumination. The YK0083 cells were exposed to blue light for 1 h in batch culture and spread on agar. The colonies were photographed under the exposure of ambient light (left) and excitation light (right). The colonies of the cells retaining the *mcherry-cat* gene were visually detectable under the excitation light (arrows). (B) Relationship between blue-light illumination duration and fractions of non-fluorescent colonies. Black points represent the means. Error bars represent standard errors (*N* = 239 for no IPTG; *N* = 327 for 0 h; *N* = 335 for 0.25 h; *N* = 373 for 0.5 h; *N* = 344 for 1 h; *N* = 350 for 2 h; *N* = 296 for 4 h; *N* = 209 for 6 h). “no IPTG” represents the condition where the cells were cultured in the M9 medium without IPTG and not exposed to blue light. (C) The correspondence between mCherry-CAT fluorescence and presence of the *cat* resistance gene. FL+ and FL-represent the presence/absence of mCherry fluorescence visually inspected for the colonies. *cat* + and *cat* - represent the presence/absence of the *cat* gene detected by colony PCR. All non-fluorescent colonies had lost the *cat* gene. All but one fluorescent colony retained the *cat* gene.

**Figure S3.**
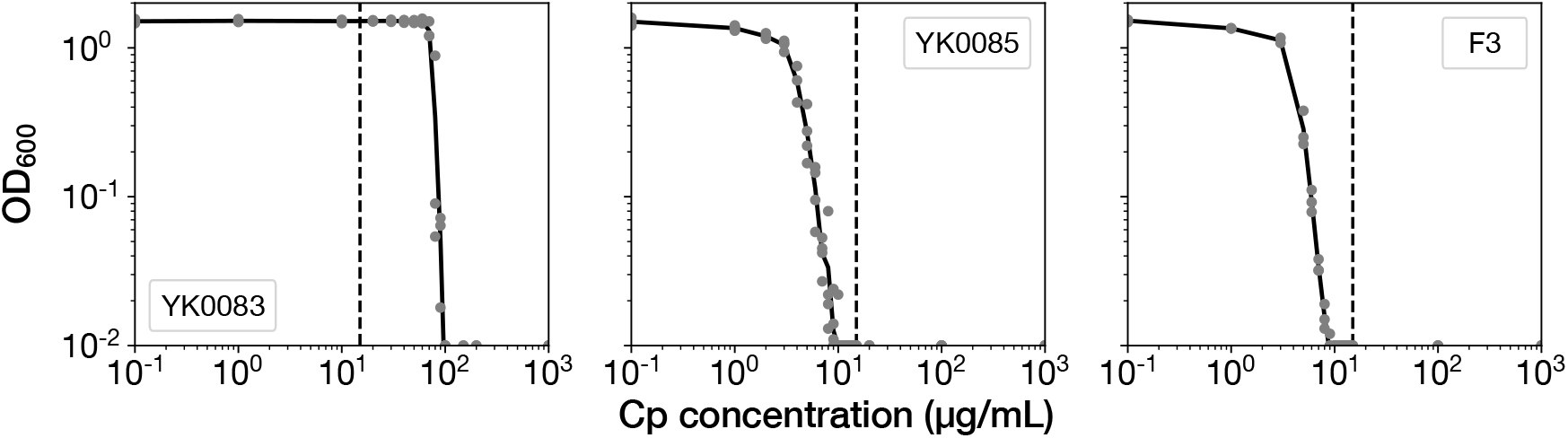
MIC tests. Gray points represent the OD_600_ of the indicated strains after a 23 h incubation period in the M9 media containing the corresponding concentrations of Cp. The minimum concentration where OD_600_ became lower than 0.01 was adopted as the MIC for each strain. 15 *µ*g/mL of Cp was used in the time-lapse experiments (dashed line). The measurements were repeated at least thrice.

**Figure S4.**
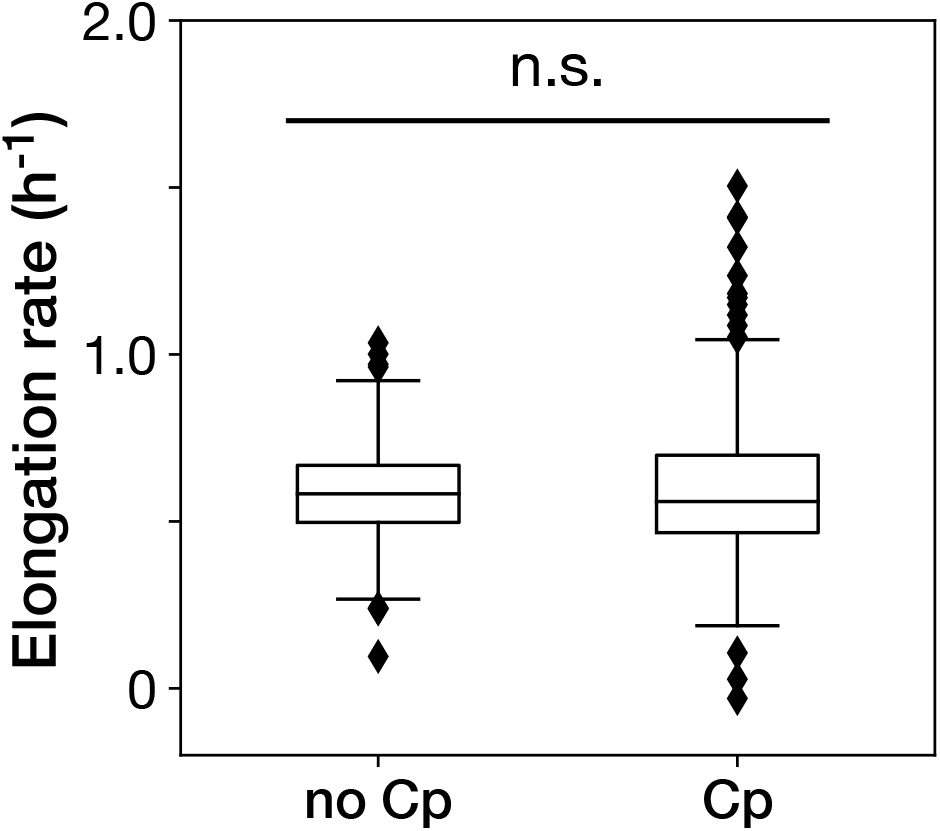
The Cp concentration used in the time-lapse measurements caused no significant effect on the growth of non-deleted YK0083 cells. Box plots show the elongation rates of the YK0083 cells growing in the mother machine. “no Cp” denotes the drug-free condition; “Cp” denotes the condition under the exposure of 15 *µ*g/mL of Cp. The elongation rates were obtained from the non-deleted cell lineages up to ten generations after blue-light illumination. We detected no significant differences in elongation rates between the two conditions (*p* = 0.63, Welch’s *t*-test).

**Figure S5.**
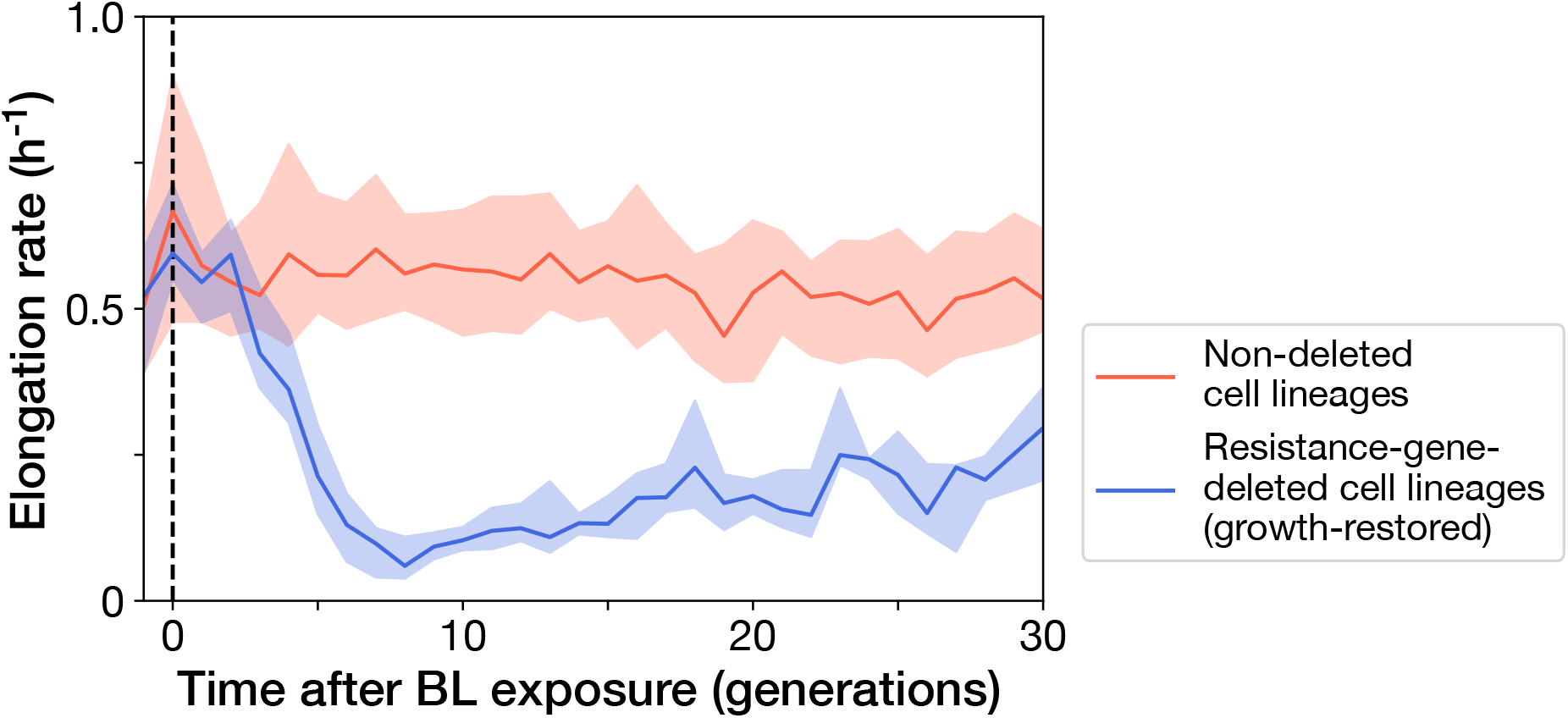
Transitions of elongation rates after blue-light illumination. The YK0083 cells were exposed to blue light for 30 min from *t* = −0.5 h to 0 h. The lines and shaded areas represent the medians and the 25-75% data ranges, respectively. Red represents the non-deleted cell lineages. Blue represents the growth-restored resistance-gene-deleted cell lineages.

**Figure S6.**
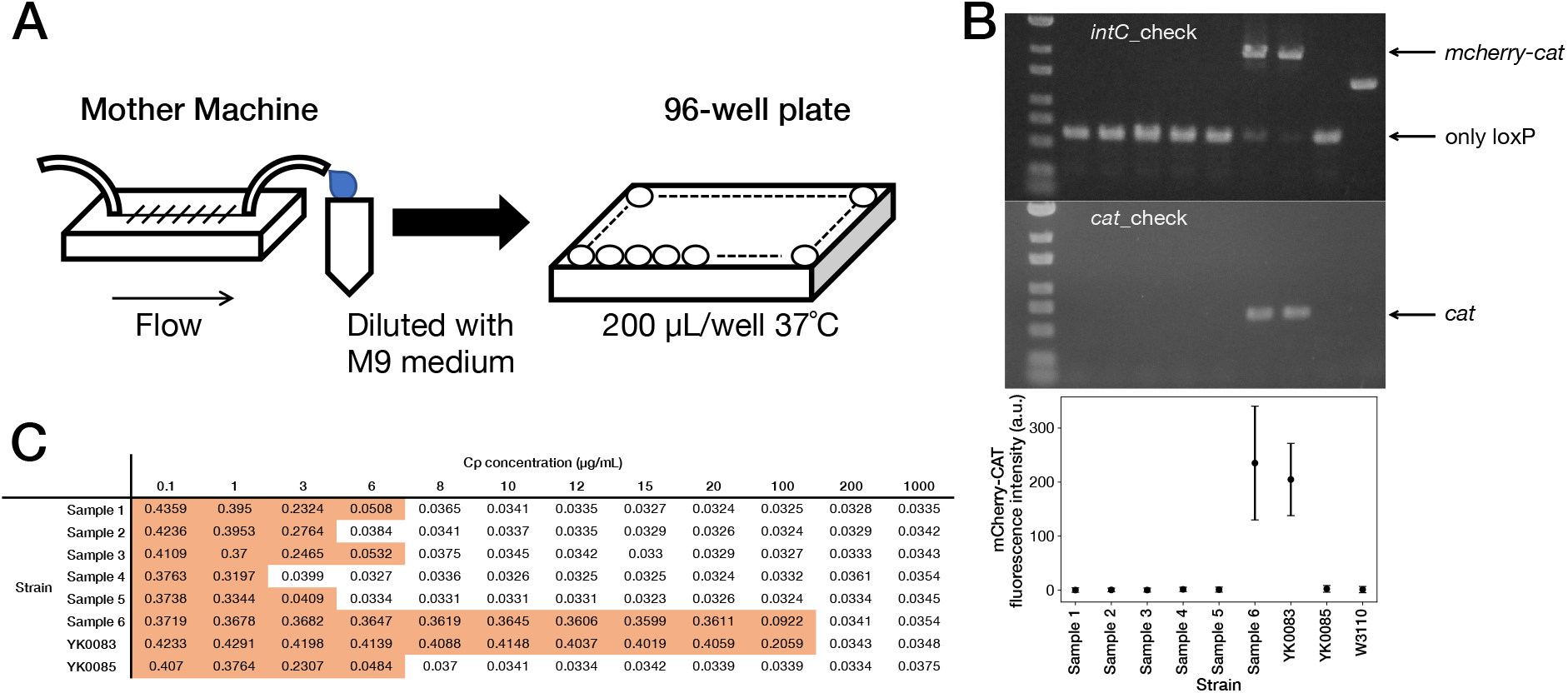
Correspondence between fluorescence loss and *mcherry-cat* gene deletion in the cells sampled from the mother machine. (A) The scheme for acquiring the resistance-gene-deleted cells from the mother machine. The culture media flowing out from the mother machine were sampled to obtain the cells that lost the *mcherry-cat* gene. The media were diluted serially, and the limited-diluted media were taken in 96-well plates and incubated. (B) Correspondence between fluorescence loss and gene deletion. The cell populations were collected from the 96-well plates, and the deletion of *mcherry-cat* gene was examined by PCR. We detected no bands of both *mcherry-cat* and *cat* for Samples 1-5. Sample 6 retained the *mcherry-cat* gene. The bottom plot shows that the absence of *mcherry-cat* gene corresponds to the loss of mCherry-CAT fluorescence. (C) MIC test. The OD_600_ values of Samples 1-6, YK0083 and YK0085 after a 23 h incubation period at the indicated Cp concentrations are shown (Start OD_600_, 0.001). The entries with colored backgrounds indicate OD_600_ > 0.04.

**Figure S7.**
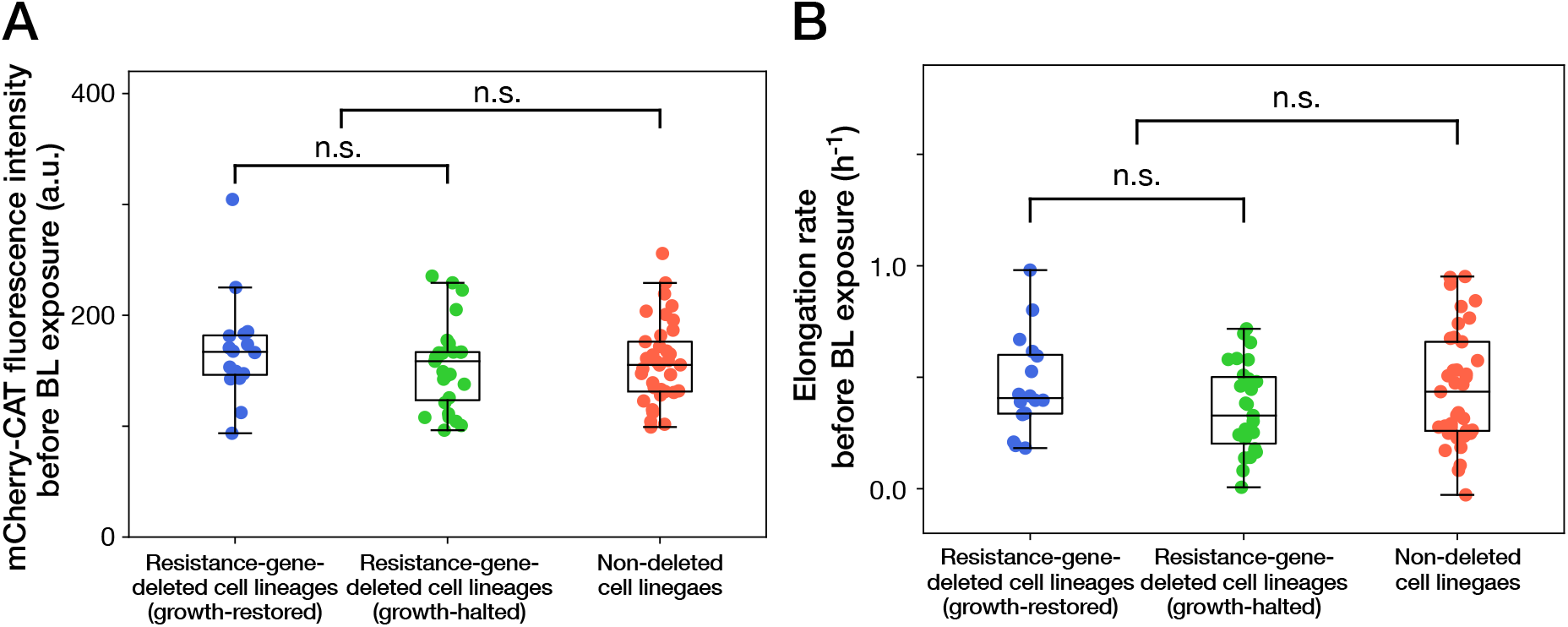
Cellular phenotypes before blue-light illumination do not correlate with the cellular fates. (A) mCherry-CAT fluorescence before blue-light illumination. Each point represents the fluorescence intensity of single-cell lineage averaged over the 1 h period immediately before blue-light illumination. Blue points represent the growth-restored resistance-gene-deleted cell lineages. Green points represent the growth-halted resistance-gene-deleted cell lineages. Red points represent the non-deleted cell lineages. *p* = 0.30 for growth-restored resistance-gene-deleted cell lineages vs growth-halted resistance-gene-deleted cell lineages; *p* = 0.85 for resistance-gene-deleted cell lineages vs non-deleted cell lineages, Welch’s t-test. (B) Elongation rates before blue-light illumination. Each point represents the elongation rate of single-cell lineage averaged over the 1 h period immediately before blue-light illumination. The color correspondence remains the same as A. *p* = 0.12 for growth-restored resistance-gene-deleted cell lineages vs growth-halted resistance-gene-deleted cell lineages; *p* = 0.35 for resistance-gene-deleted cell lineages vs non-deleted cell lineages, Welch’s t-test.

**Figure S8.**
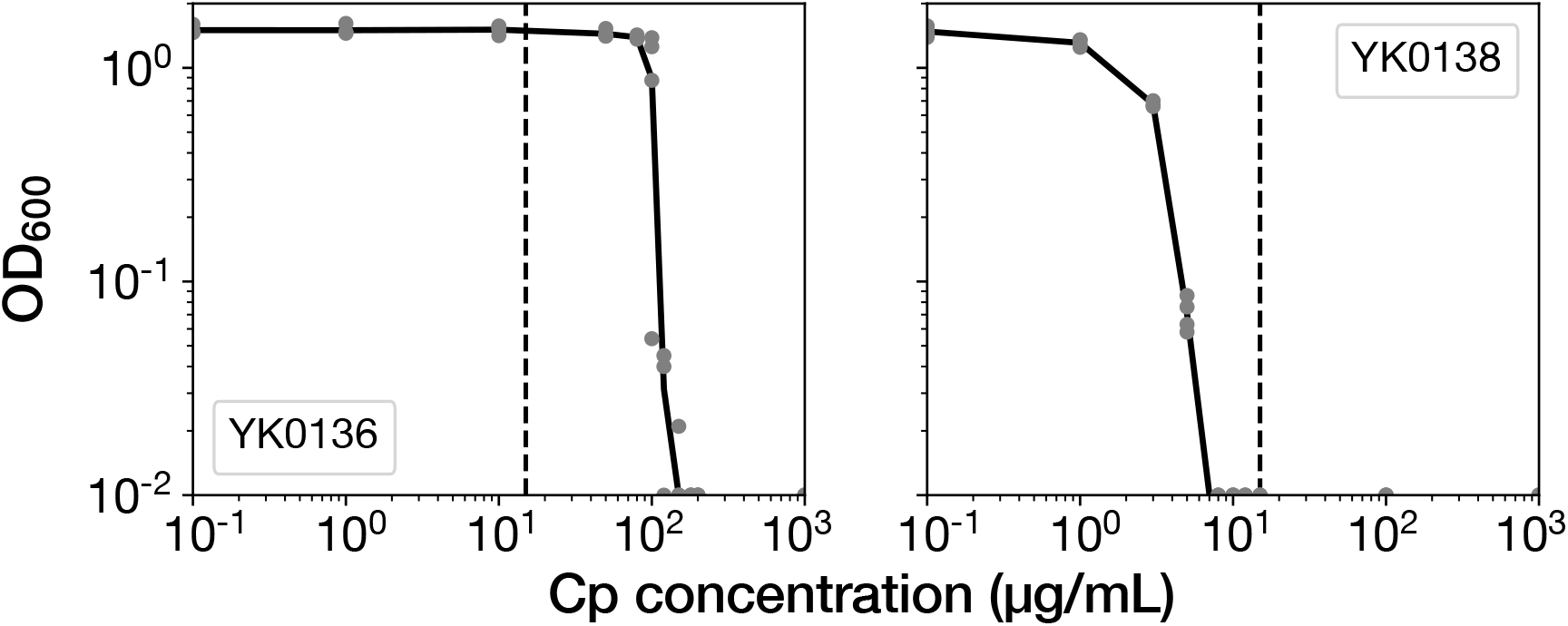
MIC tests of the ribosome reporter strains. Gray points indicate the OD_600_ of the cell cultures after a 23 h incubation period in the media containing the corresponding concentrations of Cp. The left shows the result of YK0136. The right shows the result of YK0138. The minimum concentration where OD_600_ became lower than 0.01 was adopted as the MIC for each strain. 15 *µ*g/mL of Cp was used in the experiments (dashed line). The measurements were repeated at least thrice.

**Figure S9.**
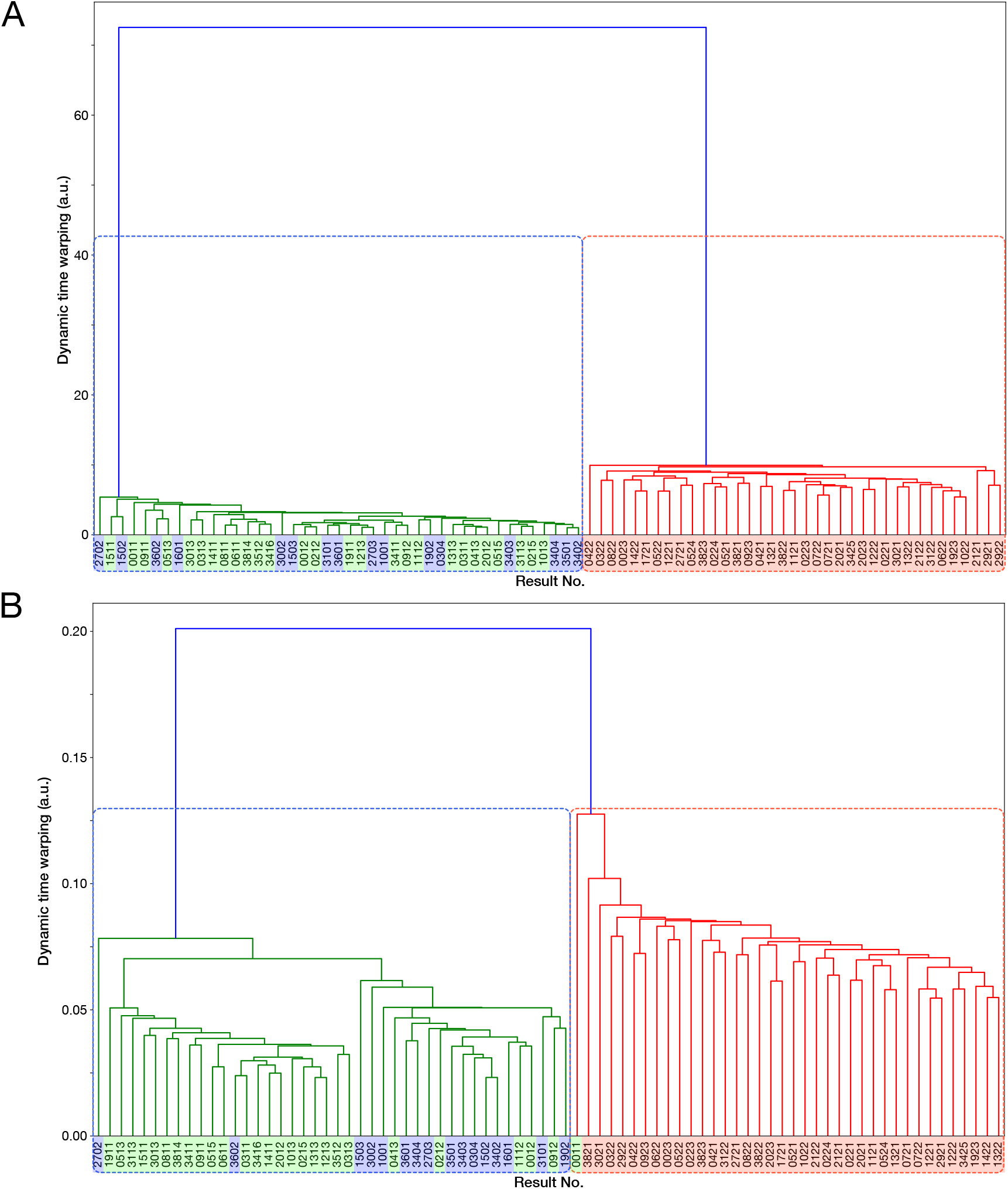
Time-course transitions of elongation rates are sufficient for classifying resistance-gene-deleted and non-deleted cell lineages. (A) Hierarchical clustering of single-cell lineages based on the transitions of mCherry-CAT fluorescence intensities. The mCherry-CAT fluorescence transitions in single-cell lineages over the entire periods of measurement were used for clustering. The distances between cell lineages were calculated by dynamic time warping. The numbers shown on the horizontal axis represent cell lineage IDs. The background colors behind the cell lineage IDs represent the fates of cell lineages assigned manually by the experimenters: Blue corresponds to growth-restored resistance-gene-deleted cell lineages; Green represents growth-halted resistance-gene-deleted cell lineages; Red represents non-deleted cell lineages. (B) Hierarchical clustering of single-cell lineages based on the transitions of elongation rates. The cell lineage clusters indicated by the blue and red dotted lines constitute two main groups, corresponding to resistance-gene-deleted and non-deleted cell lineages, respectively. Only one resistance-gene-deleted cell lineage (0011) was assigned wrongly as a non-deleted cell lineage. Therefore, the transitions of elongation rates are sufficient for classifying the resistance-gene-deleted and non-deleted cell lineages at high precision.

**Figure S10.**
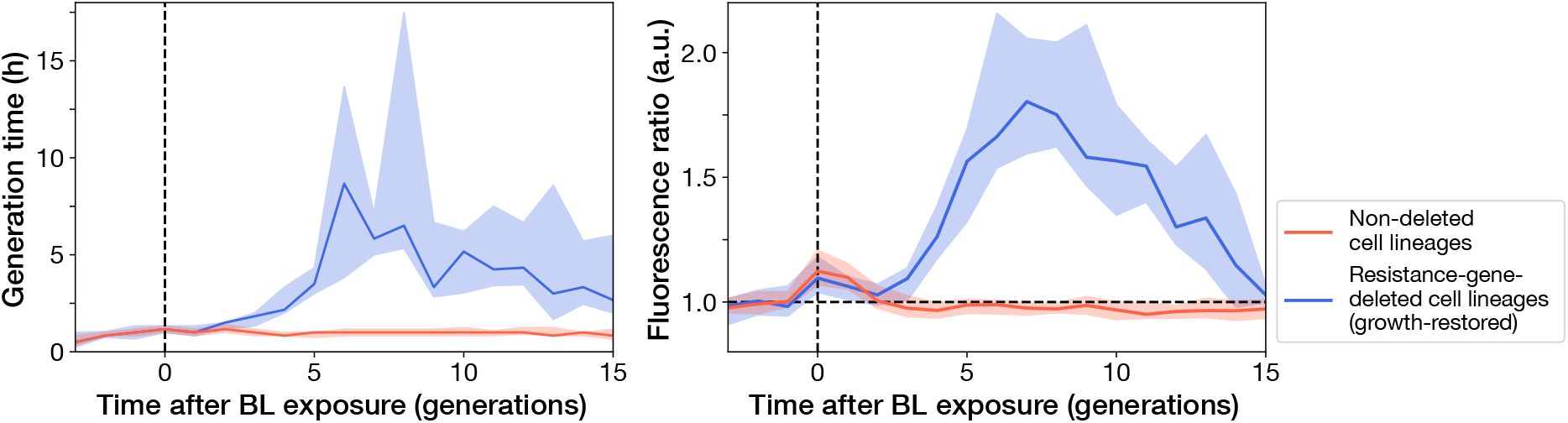
Transitions of generation time and RplS-mCherry/RpsB-mVenus fluorescence ratio observed in ribosome reporter strain shown in the time unit of generation. Left: Generation time transitions; Right: RplS-mCherry/RpsB-mVenus fluorescence ratio transitions. The lines and shaded areas represent the medians and the 25-75% data ranges, respectively. Blue corresponds to growth-restored resistance-gene-deleted cell lineages. Red corresponds to non-deleted cell lineages.

**Figure S11.**
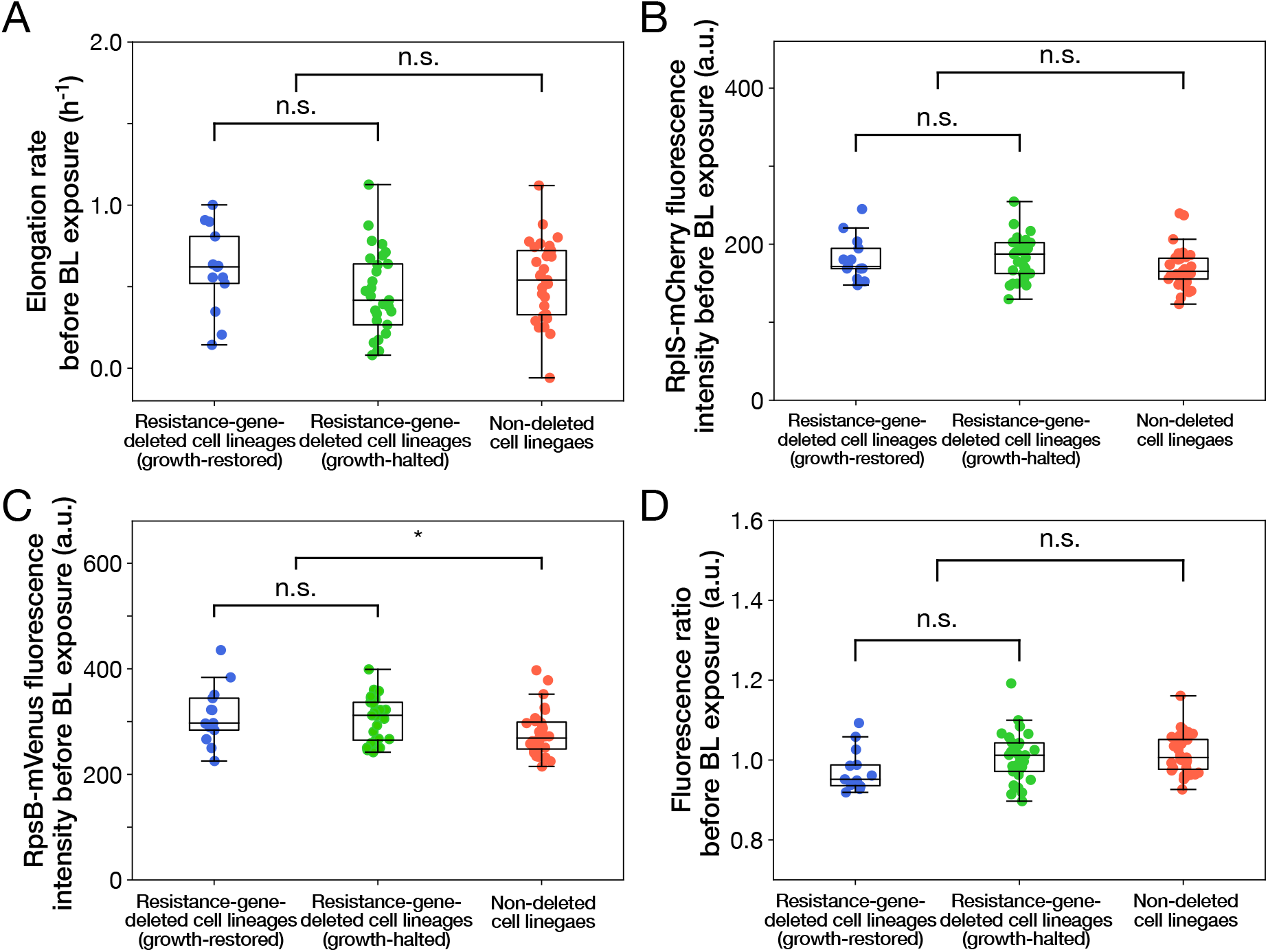
Relationships between cellular phenotypes before blue-light illumination and cellular fates. (A) Relationship between elongation rates before blue-light illumination and cellular fates. Blue represents growth-restored resistance-gene-deleted cell lineages. Green represents growth-halted resistance-gene-deleted cell lineages. Red represents non-deleted cell lineages (Color correspondence remains the same in the following figures). Each point represents the elongation rate of single-cell lineage averaged over the 1 h period immediately before blue-light illumination. *p* = 0.10 for growth-restored vs growth-halted cell lineages; *p* = 0.54 for resistance-gene-deleted vs non-deleted cell lineages, Welch’s t-test. (B) Relationship between RplS-mCherry fluorescence intensities averaged over the 1 h period before blue-light illumination and cellular fates. *p* = 0.87 for growth-restored vs growth-halted cell lineages; *p* = 0.038 for resistance-gene-deleted vs non-deleted cell lineages, Welch’s t-test. (C) Relationship between RpsB-mVenus fluorescence intensities averaged over the 1 h period before blue-light illumination and cellular fates. *p* = 0.60 for growth-restored vs growth-halted cell lineages; *p* = 0.010 for resistance-gene-deleted vs non-deleted cell lineages, Welch’s t-test. Asterisk denotes the statistical difference judged at the significance level of 0.01. (D) Relationship between RplS-mCherry/RpsB-mVenus fluorescence ratio averaged over the 1 h period before blue-light illumination and cellular fates. *p* = 0.098 for growth-restored vs growth-halted cell lineages; *p* = 0.15 for resistance-gene-deleted vs non-deleted cell lineages, Welch’s t-test.

## Notes

### Competing Interest Statement

The authors have declared no competing interest.

